# Conditioner application improves bedding quality and bacterial composition with potential beneficial impacts for dairy cow’s health

**DOI:** 10.1101/2023.12.13.571496

**Authors:** Lysiane Duniere, Bastien Frayssinet, Caroline Achard, Eric Chevaux, Julia Plateau

## Abstract

Recycled Manure Solid is used as bedding material in cow housing but can be at risk for pathogens development. Cows spend several hours per day lying, contributing to the transfer of potential mastitis pathogens from the bedding to the udder. The effect of a bacterial conditioner (Manure Pro, MP) application was studied on RMS-bedding and milk qualities and on animal health.

MP product was applied on bedding once a week for 3 months. Bedding and teat skin samples were collected from Control and MP groups at D01, D51 and D90 and analyzed through 16S rDNA amplicon sequencing. MP application modified bacterial profiles and diversity. Control bedding was significantly associated with potential mastitis pathogens while no taxa of potential health risk was significantly detected in MP beddings. Functional prediction identified enrichment of metabolic pathways of agronomic interest in MP beddings. Significant associations with potential mastitis pathogens were mainly observed in Control teat skin samples. Finally, significantly better hygiene and lower Somatic Cell Counts in milk were observed for cows from MP group while no group impact was observed on milk quality and microflora. No dissemination of MP strains was observed from bedding to teats or milk.

**Importance:** The use of MP conditioner improved RMS-bedding quality and this higher sanitary condition had further impacts on dairy cows’ health with less potential mastitis pathogens significantly associated to bedding and teat skin samples of animals from MP group. The animals also presented an improved inflammation status, while milk quality was not modified. The use of MP conditioner on bedding may be of interest in controlling the risk of mastitis onset for dairy cows and further associated costs.

## Introduction

Bedding materials can be inorganic (sand) or organic such as sawdust or wood products. Recycled manure solid (RMS) is a type of organic bedding made from fresh manure, brought to at least 34% DM through removal of water using a screw, a roller press, or high-performance slurry separation equipment. Despite this process, several pathogens have been isolated from RMS bedding (1). For example, higher detection of *Salmonella* spp and *Listeria monocytogenes* was observed in unused RMS compared to unused straw beddings (2). Due to high DM content sand and sawdust are often considered as “safer” bedding material, however pathogens such as coliforms, *Klebsiella* and *Streptococcus* were found in both bedding materials in free-stall dairy farm (3). Clean stall bedding is very closely related to animal hygiene and bacterial exposure. Dairy cows lying time has been measured as 10h to 13h per day (4) and during this period, cows’ teats are in direct contact with bedding materials. Dirty bedding, associated with poor animal hygiene and soiled udder, has been recognized as a source of intramammary infections and an increasing risk of mastitis onset (5–8). The main pathogens responsible for mastitis are *Streptococcus*, *Staphylococcus*, *Escherichia*, *Klebsiella* or *Corynebacterium*, but up to 24 bacterial genera have been identified from udder pathogenic isolates (9,10). Mastitis is classified as clinical or subclinical depending on the visibility of mammary gland inflammation effects. Subclinical mastitis does not produce visible effects on milk quality but has important impacts on milk composition, mainly through an increase in somatic cell counts (SCC), while clinical mastitis is associated with abnormal milk detection of flakes and clots and visible pathological changes of the mammary tissue (red and swollen udder, fever…) (11). Bovine mastitis has thus a massive negative effect on animal well-being and also on farm economics due to treatment costs, reduction in milk production and early culling (12–17). It is one of the most frequent diseases in dairy industry and in a meta-analysis gathering 372 studies spanning the period 1967-2019, sub-clinical mastitis prevalence was observed at 42 % worldwide while clinical mastitis reached 15 % prevalence (18).

Results are unclear about the specific impact of RMS bedding on mastitis occurrence in dairy cows. RMS was suggested to be associated with lower udder health (19) and increased sub-clinical mastitis risks (20) compared to other organic or inorganic beddings, while no correlation was found between RMS bedding and sub-clinical mastitis (21), animal hygiene (22) and SCC in milk (23). Only few studies were published on the interest of bedding conditioners regarding mastitis prevention. Chemical disinfectants can be used to limit bacterial counts in organic bedding and decrease their transfer on teat skin (24,25) but the effects are variable depending on bedding material, pathogens considered and duration of antibacterial activity (26,27). Moreover, some bedding conditioners such as hydrated lime (calcium hydroxide, Ca(OH)_2_) are caustic and can cause animal and farm worker’s skin damage. The use of specific bacteria meant to drive the microbial populations through fermentation is widely used in agronomy (silage inoculant (28), composting process (29), …). Bioremediation and ecological niche occupation with specific bacterial strains have also been successfully employed in soil depollution (30) or positive biofilm production in food industry for example (31,32). The bedding conditioner tested in this study is composed of a cocktail of *Bacillus* and *Pediococcus* strains and enzymes aiming at promoting beneficial bacteria and limiting the presence of undesirable microorganisms through the development of a biofilm. *Bacillus* are producing a great variety of extra-cellular enzymes such as α-amylase, cellulase and proteases involved in organic matter degradation (33). However, they are recognized as milk spoilage bacteria causing shelf-stability issues in dairy products including ropiness, sweet curdling and off-flavours (34) and their presence or potential transfer in milk should be avoided carefully.

Providing clean and safe standing and lying environment by controlling bedding microbial quality is thus a key step in mastitis control. Up to now, no study on the effects of bacterial bedding conditioner on dairy cows’ bedding quality and further animal health has ever been published. The aim of this study was 1) to study the effect of a bacterial bedding conditioner (Manure Pro®, Lallemand SAS, France) on RMS-bedding physico-chemical parameters and bacterial populations, 2) to evaluate the impacts of this bedding treatment on animal hygiene, health and teat skin bacterial populations and finally 3) to assess the absence of negative impact on milk.

## Results

### Physico-chemical parameters and bacterial population of bedding samples

The evolution between the 2 groups tended to differ between D01 and D51 (Period 1) with a decrease of DM observed in Control (-10.39 % DM) while DM of treated bedding slightly increased (+1.97 % DM, Table 1). A tendency for a stronger pH increase was also noted for MP bedding during the same period (+0.20 and +0.55 pH unit for Control and MP, respectively). No significant evolution of the measured parameters was observed in Period 2.

**Table 1:** Physico-chemical parameters of bedding samples measured at D01, D51 and D90 in Control and MP group (n=6). Mean, SEM and p-values associated during period 1 (D0-D51) and Period 2 (D51-D90).

|  | Control |  |  | MP |  |  |  | p-values |  |
| --- | --- | --- | --- | --- | --- | --- | --- | --- | --- |
|  | D01 | D51 | D90 | D01 | D51 | D90 | SEM | Period 1 | Period 2 |
| DM | 61.01 | 50.62 | 54.83 | 57.61 | 59.58 | 50.71 | 1.199 | 0.053 | 0.130 |
| pH | 8.76 | 8.96 | 9.07 | 8.52 | 9.07 | 9.09 | 0.038 | 0.098 | 0.247 |
| OM | 39.33 | / | 34.84 | 33.76 | / | 37.84 | 1.258 | 0.250 |  |

Both experimental bedding groups were characterized by stable relative abundance of Firmicutes (average of 33.03 ± 7.72%) and Proteobacteria (average of 26.75 ± 5.16%) over time and a transient decrease of Actinobacteria at D51 (Suppl Figure 1A). Families associated to potential and recognized mastitis pathogens (ie Staphylococcaceae and Streptococcaceae) were identified in both Control and Manure Pro groups (Suppl Fig 1B).

Bacterial richness didn’t vary significantly according to Time but was close to significance for Group (Observed Amplicon Sequence Variants (ASVs), p=0.053) with higher richness observed in MP samples (Figure 1A). A transient decrease of evenness was observed at 51 days in both Control and MP groups (Shannon, Time effect, p=0.04) (Figure 1B). A significant Group effect was also observed (p = 0.009) with greater Shannon value for MP compared to Control samples highlighting the higher heterogeneity of bacterial populations in MP samples at day 90. A clear evolution of bacterial profiles was observed at ASV level in all bedding samples over time (Figure 1C, p= 0.001) while a Group effect could be observed only after 90 days (p=0.003) between Control and MP samples.

**Figure 1:**
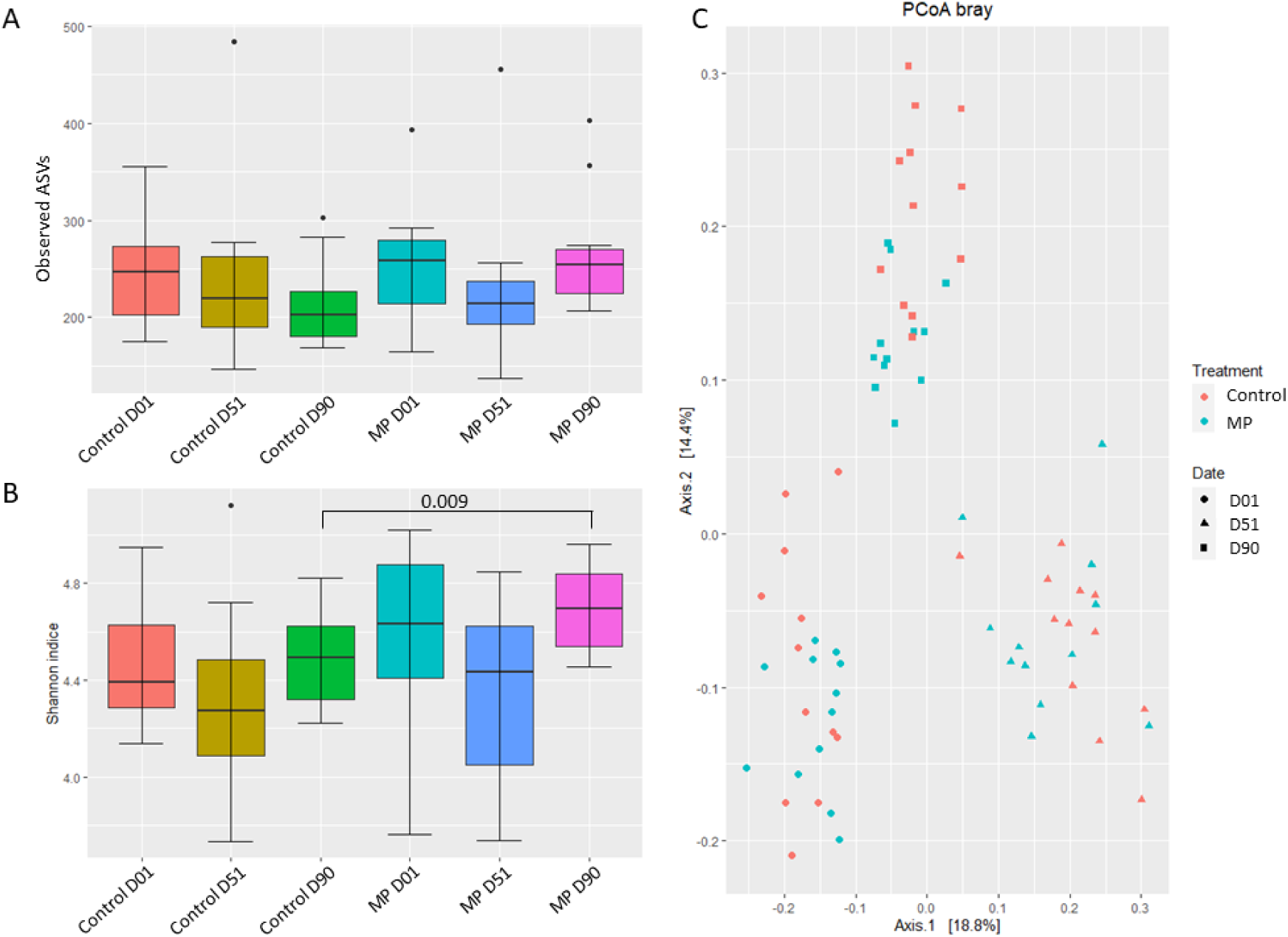
alpha diversity indices expressed through A) Observed ASVs and B) Shannon index and beta diversity expressed through C) PCoA plot based on Bray–Curtis distance for normalized abundance data of Control and MP samples at D01 D51 and D90 (n=12).

Differential analysis was performed with Indicspecies R package which determines lists of ASVs that are more specifically associated to each experimental group through an Indicator Value (IndVal) index. IndVal index is a combination of A and B components. Component ‘A’ represents the *specificity* or *positive predictive value* of the species as indicator of the target experimental group. A specie with A=1 is a good indicator of the experimental group as it is found only in this specific group. Component ‘B’ is called the *fidelity* or *sensitivity* of the species as indicator of the target experimental group. A B=1 specie indicates that this specie is observed in all samples from the experimental group considered. In order to circumvent the issue of taxonomic multi-affiliation occurring for several species belonging to *Streptococcus* or *Staphylococcus* genera, decision was made to agglomerate ASVs obtained from bioinformatic analysis up to genus level. Linear discriminant analysis (LDA) Effect Size (LEfSe) analysis, commonly employed in metagenomic ecological studies, was used as a complementary differential analysis to confirm the taxa identified through IndicSpecies analysis.

Similar numbers of taxa were identified as characteristics of each experimental group with highest and lowest number of taxa identified in MP at D01 and D51 (36 and 12 taxa identified respectively) (Table 2). Interestingly, taxa characteristics of MP groups were identified as environmental nonpathogenic bacteria such as *Mesorhizobium*, Xanthomonadaceae family or commensal digestive bacteria such as Lachnospiraceae NK3A20 group and Christensenellaceae R7 group and no taxa linked to potential pathogen was identified (Supplementary Material).

**Table 2:**
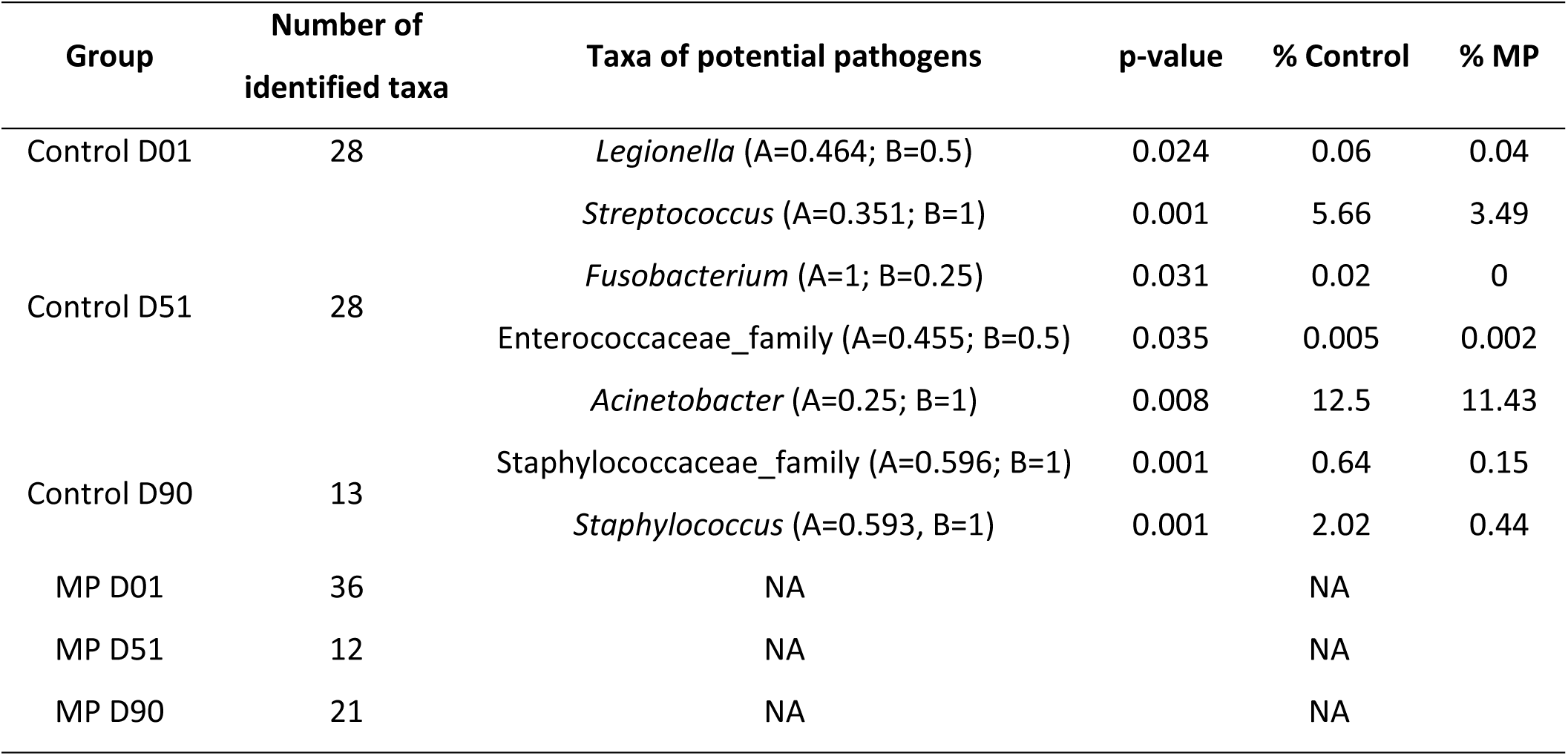
Number of taxa and taxa of potential pathogens identified through Indicspecies analysis, p-values associated and relative abundances observed for Control and MP beddings analyzed at D01, D51 and D90.

On the contrary, taxa belonging to genera responsible for mastitis were associated to Control groups at different sampling times. *Streptococcus* and *Acinetobacter* were identified at D51 and a member from Staphylococcaceae family and *Staphylococcus* were also observed at D90 in all samples of the Control group of interest (B = 1). Other potential pathogens were associated to Control group at D01 or D51: *Fusobacterium* was only observed in D51 samples (A = 1) as well as a member of Enterococcaceae family while *Legionella* was significantly associated to D01. Similar results were obtained through LEfSe analysis (Figure 2) as *Acinetobacter* and *Streptococcus* were associated to Control D51 and *Staphylococcus* to Control D90 while the only taxa associated to MP treatment was the non-pathogenic genus *Flavobacterium* at D51. Noteworthy, 2 ASVs assigned to Bacillaceae family and *Bacillus* genus were associated with Control D51 through Indicspecies analysis (A=0.48, B=0.75 and A=0.35, B=0.92 respectively) while none was associated to MP groups.

**Figure 2:**
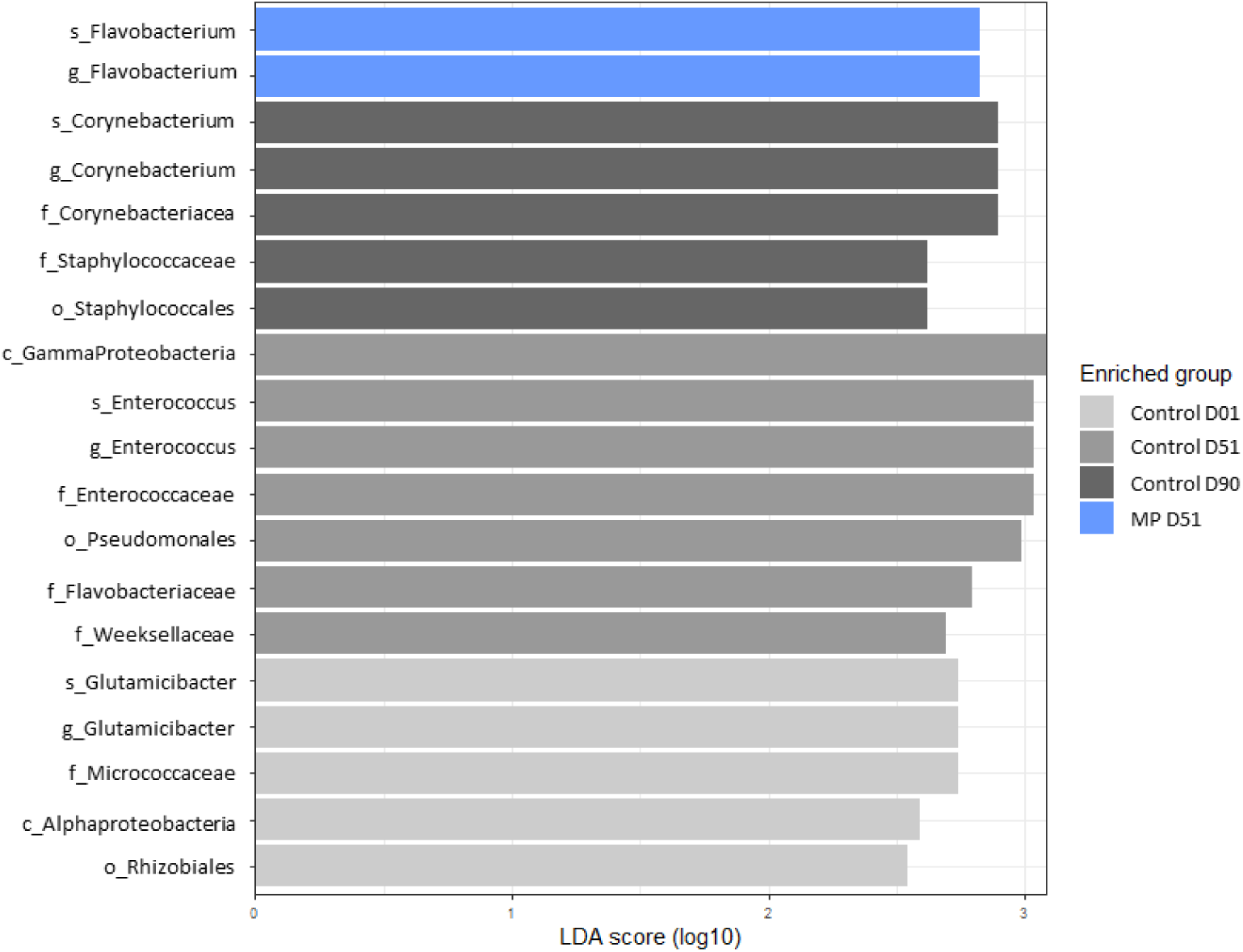
LEfSe analysis identifying specific taxa agglomerated to genus level on Control (grey bars) and MP (blue bars) samples at D01, D51 and D90 (LDA cuttoff 2.5, p < 0.05).

The PICRUSt2 pipeline was used to implement a functional prediction for the DADA2 filtered reads using MetaCyc pathway database. The comparisons between Control and MP samples considering each sampling time separately identified few significantly different potential metabolic pathways (Figure 3, p < 0.05). No significant difference was observed between Control and MP at D01 while 2 pathways linked to biogenic amines production were predicted to be more abundant in Control than in MP samples at D51 and 7 pathways linked to several biological functions, were identified in higher proportions in MP group at D90, notably a pathway of nitrifier denitrification which might be of interest in bedding management.

**Figure 3:**
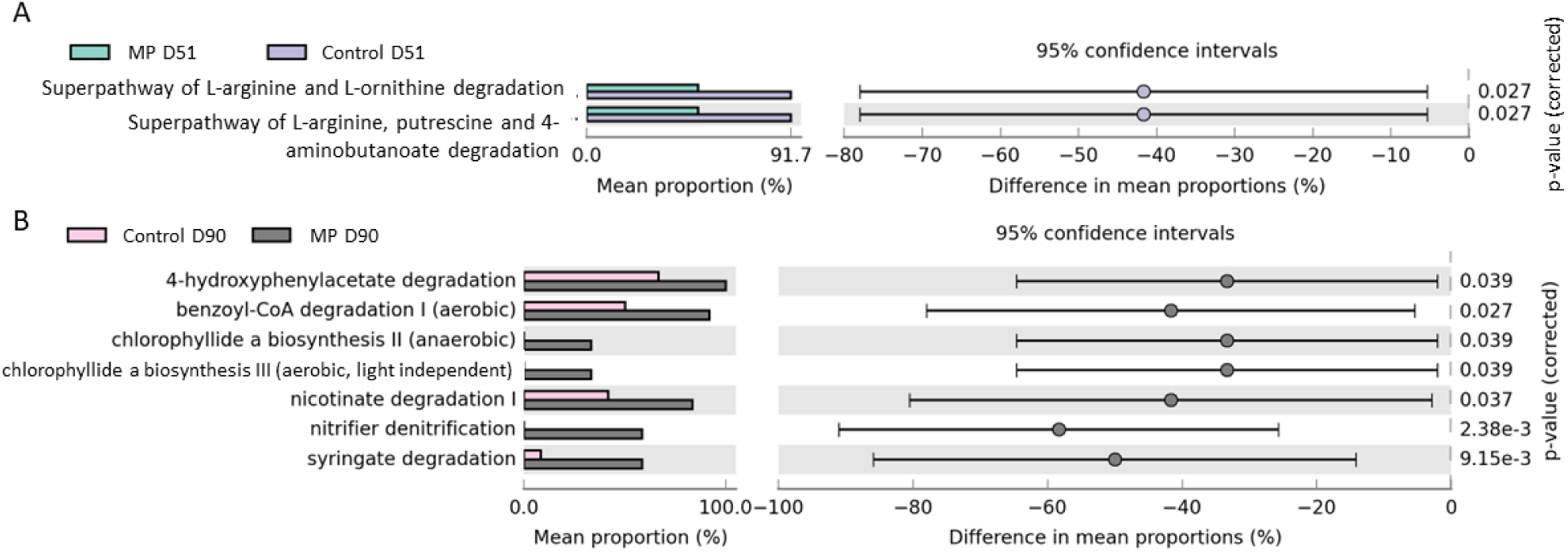
Statistically different metabolic pathways identified in Control and MP samples at A) D51 and B) D90 identified through PICRUSt2 pipeline prediction

### Animal health and teat skin bacterial population

Animal hygiene was assessed through cleanliness score, concentration of teat skin culturable microflora (Table 3) and SCC in milks over the trial (Figure 4). A significant Time by Group interaction was observed for these 3 parameters. At D51, a significantly deteriorated hygiene was observed concomitantly to a numerically higher microbial concentration on teat skin of Control cows compared to MP animals.

**Figure 4:**
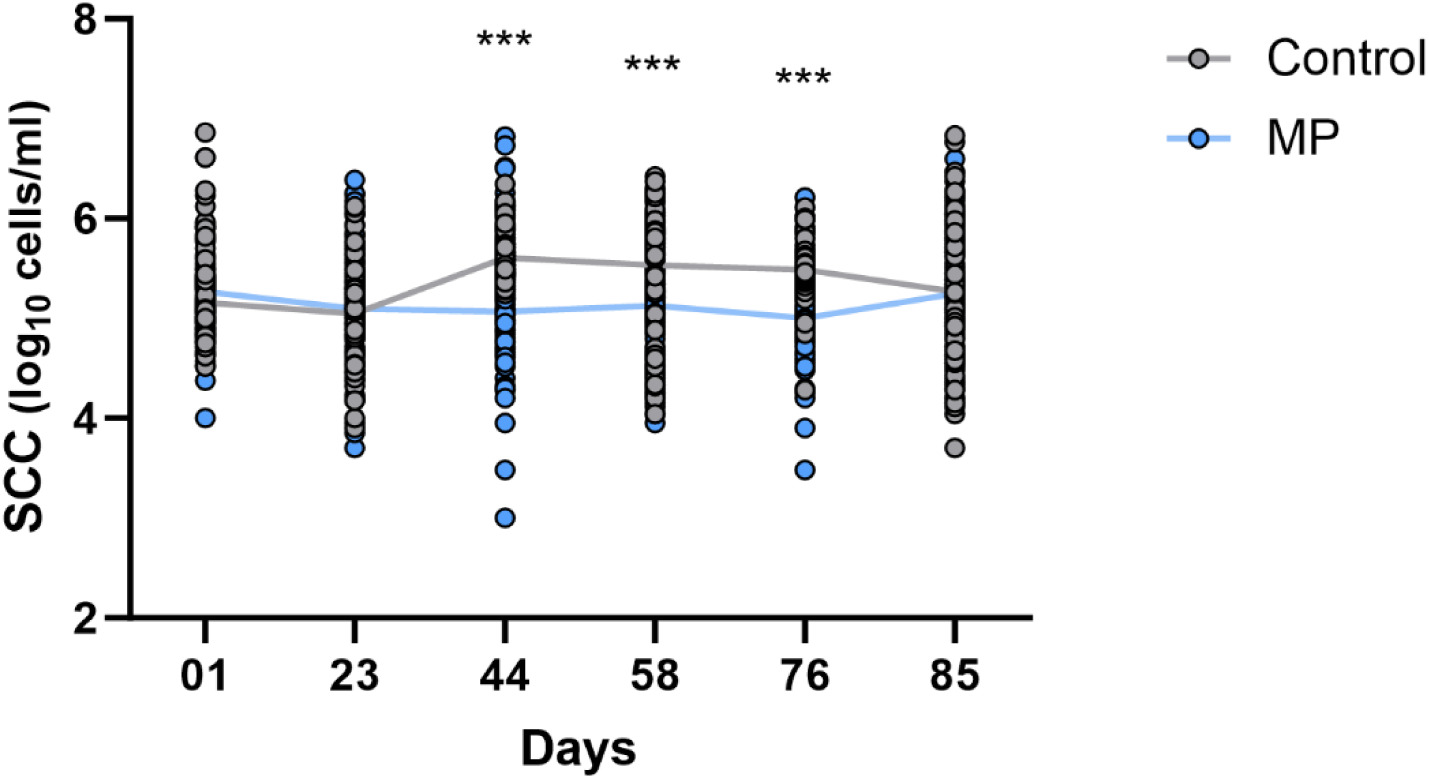
Log_10_ SCC/ml in milk collected from cows assigned to Control (Grey, n=113) or MP (Blue, n=115) group over the trial. *** indicates a post-hoc Sidak’s pvalue < 0.0001

**Table 3:**
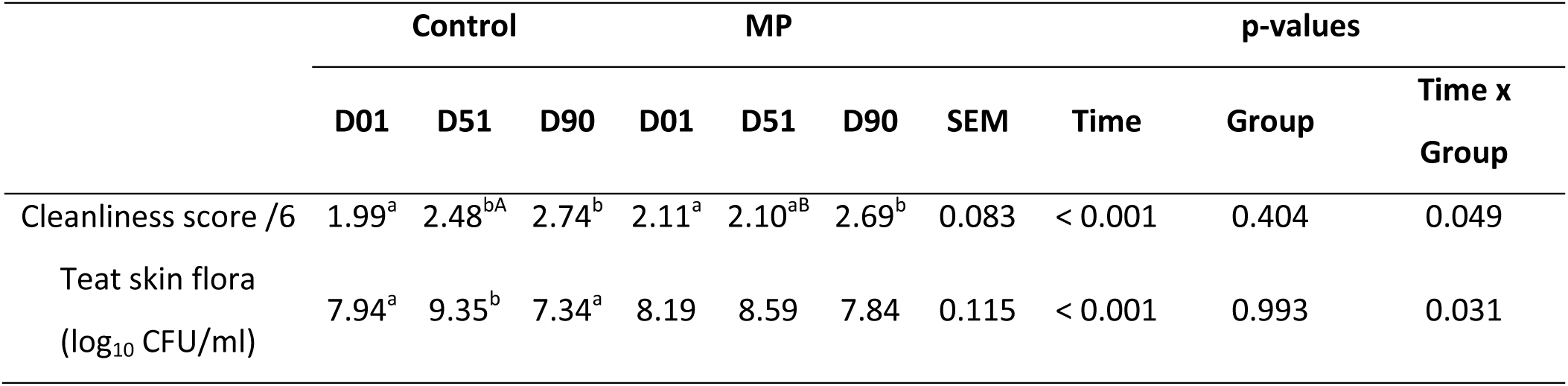
Animal hygiene parameters (cleanliness score and concentration of teat skin flora) observed in Control and MP cows at D01, D51 and D90 (n=20). Mean, SEM and p-values associated. Lower superscript letters indicates a significant difference across time and capital superscript letters a significant difference between groups.

SCC in milks was strongly decreased in MP group with significantly lower somatic cell concentration between D44 and D76 compared to milk from Control cows (Figure 4). A highly significant effect of both Time and Group factors on SCC was also observed over the trial (p < 0.0001). Considering a 250 000 cells/ml threshold, 305 milk samples from cows in Control group have been measured above this limit during the trial for only 151 milk samples in MP group (Fisher test, p < 0.0001).

Finally, the number of mastitis suspicions based on milk conductivity above 90 mS/cm was not significantly different between Control and MP groups (p > 0.05, Suppl Table 1) either when cases were considered per week or over the entire experimental period. Although not significant, the total number of mastitis suspicion cases leading to an effective pathogen identification through on-farm culture plating was higher in Control than in MP group (57 cases in Control and 51 cases in MP group respectively, p=0.66), with a higher number of suspicion cases identified as environmental mastitis pathogens (35 cases in Control and 22 cases in MP group respectively, p=0.143).

As previously observed for bedding bacterial composition, the relative abundances of bacterial phyla in teat skin samples were quite stable over time in both Control and MP groups (Suppl Fig 2A). Teat skin bacterial community was dominated by Firmicutes (38.76 ± 12.30%), followed by Proteobacteria (22.79 ± 7.79 %), Bacteroidota (20.38 ± 7.39 %) and Actinobacteria (17.45 ± 7.13 %). At Family level, similar profiles were observed between Control and MP samples and an evolution over time was observed at D90 with an increase in Pseudomonadaceae (from 2.335 % to 6.44 %) and a decrease in Enterococcaceae (from 6.06 % to 2.13 %) and Streptococcaceae (from 4.10 % to 1.16 %) families (Suppl Fig 2B). Bacterial families including potential and recognized mastitis pathogens such as Streptococcaceae (3.21 ± 4.57 %) and Staphylococcaceae (2.55 ± 2.18 %) were observed in Control and MP groups with variations over time. A significant effect of Time (Observed ASVs, p < 0.001) and a trend for Group effect (Shannon, p = 0.073) was observed on alpha diversity indices, highlighting a transient decrease of bacterial diversity at D51 and an overall higher diversity of bacterial teat population in MP group (Figure 5A and 5B). The temporal evolution of bacterial profiles of both Control and MP samples was also observed through beta diversity analysis (Figure 5C, p < 0.001 for each sampling time). Noteworthy, a significant difference was observed between MP and Control at each sampling time (p_D01_=0.008, p_D51_=0.033 and p_D90_=0.003).

**Figure 5:**
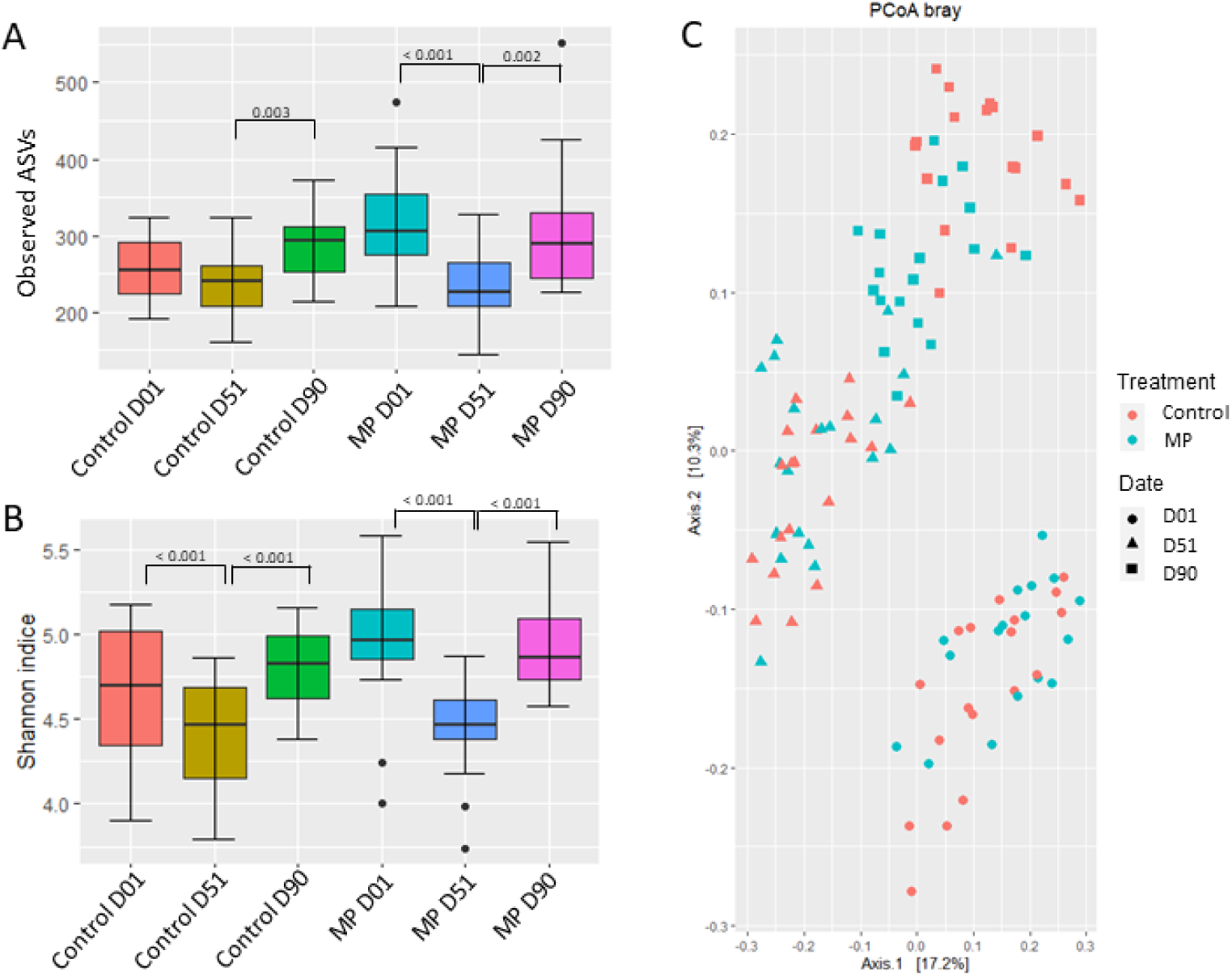
alpha diversity indices expressed through A) Observed ASVs and B) Shannon index and beta diversity expressed through C) PCoA plot based on Bray–Curtis distance for normalized abundance data of Control and MP teat samples at D01 D51 and D90 (n=18).

Indicspecies analysis identified more taxa characteristics of MP samples compared to Control samples at both D01 and D90 (Table 4). Interestingly, lactic acid bacteria were significantly associated to Control samples at D01 and *Bifidobacterium* at D90. Potential mastitis pathogens such as *Trueperella* and a member of Staphylococcaceae family were observed in almost all Control samples at D90 and a member of the Corynebacteriaceae family was associated to Control D01 samples. Although the mean relative abundance was higher in Control group (11.33%), MP samples were specifically associated with *Acinetobacter* at D51 as all those samples harbored this genus (B=1). *Pseudomonas* was also particularly observed in MP samples at D90. No taxa associated to potential pathogens was observed in MP sample at D01 nor in Control samples at D51. Interestingly, the same 2 ASVs of Bacillaceae family and *Bacillus* associated to Control D51 in bedding samples were also observed in Control teat samples at D51 (A=0.62, B=0.84 and A=0.54, B=0.95 respectively).

**Table 4:** Number of taxa and taxa of potential pathogens identified through Indicspecies analysis, p-values associated and relative abundances observed for Control and MP teats analyzed at D01, D51 and D90.

| Group | Number of identified taxa | Taxa of potential pathogens | p-value | % Control | % MP |
| --- | --- | --- | --- | --- | --- |
| Control D01 | 11 | *Lactococcus* (A=0.47; B=1) | 0.001 | 2.15 | 0.41 |
|  |  | *Latilactobacillus* (A=0.80; B=0.55) | 0.001 | 0.14 | 0.02 |
|  |  | *Secundilactobacillus* (A=0.66; B=0.44) | 0.002 | 0.13 | 0.02 |
|  |  | Corynebacteriaceae family (A=0.55, B=0.33) | 0.009 | 0.004 | 0.0009 |
| Control D51 | 26 | NA |  | NA |  |
| Control D90 | 47 | *Bifidobacterium* (A=0.31; B=1) | 0.006 | 1.18 | 0.43 |
|  |  | *Trueperella* (A=0.33; B=0.94) | 0.002 | 0.18 | 0.07 |
|  |  | Staphylococcaceae family (A=0.29; B=0.94) | 0.001 | 0.17 | 0.13 |
| MP D01 | 76 | NA |  | NA |  |
| MP D51 | 17 | *Acinetobacter* (A=0.27; B=1) | 0.001 | 11.33 | 10.99 |
| MP D90 | 54 | *Pseudomonas* (A=0.31, B=1) | 0.001 | 5.01 | 7.34 |

LEfSe analysis significantly associated an unknwon genus of Corynebacteriales order and *Streptococcus* to Control D01 and D51 teat samples while MP D51 was associated to an unknwon genus of Moraxellaceae, a non-mastitis pathogen family (Figure 6).

**Figure 6:**
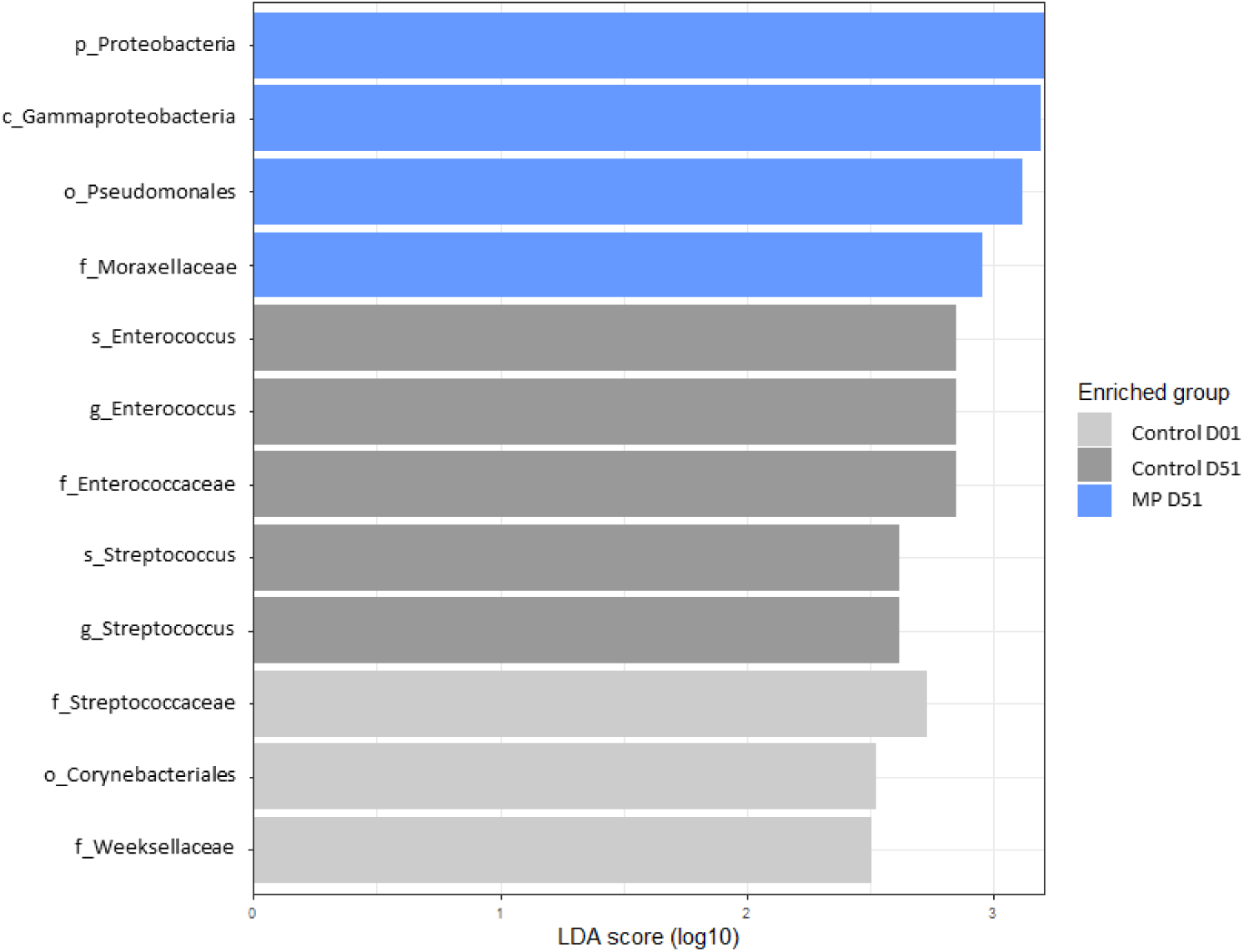
LEfSe analysis identifying specific taxa agglomerated to genus level on Control (grey bars) and MP (blue bars) samples at D01, D51 and D90 (LDA cutoff 2.5, p < 0.05).

### Milk production and quality

Milk production was followed for all enrolled animals in Control and MP groups by adding the quantity produced at each milking per day for each animal and averaged by month of treatment application (Month 1: D01-D30, Month 2: D31-D60; Month 3: D61-D90) (Suppl Fig 3). No significant difference was observed for Time, Treatment or the interaction of the 2 factors.

Milk quality was assessed through total culturable microflora and aerobic spore concentration (Table 5). No significant effect of Group was observed on these parameters over time. The total milk flora increased up to 4.3 log_10_CFU/ml at D90 in Control group while it reached its highest concentration (4.51 log_10_ CFU/ml) at D51 in MP. Spore counts were ranging between 10 and 14.06 spores/ml on average in Control group and between 12.65 and 15.59 in MP group.

**Table 5:** Total flora (log_10_ CFU/ml) and aerobic spores (counts/ml) measured in milk from Control or MP groups at D01, D51 and D90 (n=20).

|  | Control |  |  | MP |  |  | SEM | Pvalues |  |  |
| --- | --- | --- | --- | --- | --- | --- | --- | --- | --- | --- |
|  | D01 | D51 | D90 | D01 | D51 | D90 |  | D01 | D51 | D90 |
| Tota flora (log<sub>10</sub> CFU/ml) | 3.94 | 4.27 | 4.30 | 4.05 | 4.51 | 4.22 | 0.061 | 0.273 | 0.256 | 0.981 |
| Aerobic spore<br>(counts/ml) | 14.06 | 10.00 | 11.18 | 14.18 | 15.59 | 12.65 | 1.973 | 0.850 | 0.850 | 0.506 |

## Discussion

The use of RMS as bedding material has been increasing over the last years as an interesting valorization strategy for cattle organic wastes (ie fresh manure). Several studies focused on the presence of potential pathogens in this environment but were mainly performed using cultural detection methods (1,23) and very little data are available on the evolution of the overall bacterial populations through amplicon sequencing (35). Moreover, the interest of bedding conditioners to improve beddings quality have been poorly studied. To our knowledge, this study is the first one assessing the effect of a cocktail of bacteria (*Bacillus velezensis*, *Pediococcus acidilactici* and *P. pentosaceus*) and enzymes on the quality of RMS-bedding with a focus on potential mastitis pathogens.

### MP application improved bedding physico-chemical and bacterial quality

The sanitary safety of RMS is based on the fact that microorganisms initially present in fresh manure will not develop or survive in the new environmental conditions prevailing in RMS (i.e. high DM and alkaline pH), thus limiting further risks for animal and human health (36). Physico-chemical values obtained for bedding in both groups were in line with literature (23,37) with MP application leading to higher DM and pH values than Control, indicating a less favorable environment for opportunistic microbial communities.

Bacterial communities were impacted by MP application with a higher diversity observed in MP beddings. It is commonly admitted that a high microbial diversity increases the robustness of a given environment after a disturbing event, as defined by its resistance, resilience and functional redundancy (38). A high microbial diversity indicates a wide ecological niches occupation and limits the possibility for allochthonous microorganisms to establish and proliferate (39). The loss of diversity observed in case of dysbiosis generally allows a parallel growth of potential pathogenic microorganisms (40). Our results agree with this observation as differential analysis significantly associated several potential pathogens with Control groups, while MP-bedding groups were mainly associated with environmental and ruminant digestive bacteria (*Mesorhizobium*, Lachnospiraceae NK3A20 …). More precisely, bacterial genera such as *Streptococcus*, *Acinetobacter* and *Staphylococcus* were identified in higher relative abundance and prevalence in Control group over the trial. In a meta-analysis, Krishnamoorthy and colleagues have notably estimated the prevalence of *Staphylococcus* and *Streptococcus* species in subclinical or clinical bovine mastitis at respectively 28 % and 12 % worldwide over the 1979-2019 period (18). In a large scale study, *Staphylococcus*, *Streptococcus* and *Acinetobacter* were identified in 39.03 %, 11.01 % and 3.38 % of the 1 153 mastitis milk samples analyzed (41). Due to technical limitation of taxonomic assignation with 16S rDNA amplicon sequencing, some genera have been identified at higher taxonomic level, such as family level (Staphylococcaceae and Enterococcaceae). Among Enterococcaceae, *Enterococcus faecalis* and *E. faecium* are known mastitis pathogens, isolated from 18 % and 1.75 % of 2 000 mastitis milks respectively (42) and *E. faecalis* abundance have been shown to dramatically increase in mastitis milks compared to normal raw milk (43). These bacterial species have been previously observed in RMS-bedding with great variations in relative abundances (35) but the average lower prevalence observed in our study in MP beddings would likely decrease the risk of bacterial transfer to cows teats and ultimately the risk of mastitis.

Other potential pathogens, not always clearly associated to mastitis, have also been associated to Control group. *Fusobacterium* is an opportunistic pathogen, causing significant health issues in cattle such as liver abscesses and foot rot (44). *Fusobacterium necrophorum* has been identified in bovine summer mastitis (45), and in a Danish study, *F. necrophorum* has been isolated among other pathogens in 52 % of the milk samples tested (46). *Legionella pneumophila* is a well-known cause of respiratory infections in humans. *Legionella* infections have been rarely documented in animals, and cattle is only considered as accidental host for Legionellae (47). The low relative abundance of *Legionella* DNA in bedding samples in our study is probably linked to environmental contamination through ground or potable water for example (48).

The mechanisms by which MP product can modify bacterial profile of bedding remain not fully understood at this stage. *B. velezensis* has been used in agricultural industry, as an alternative to chemical fertilizers or pesticides, for its capacity to degrade toxic by-products or lignocellulose, and even for its bio-emulsifying properties (33). The functional inference analysis identified the enrichment of nitrifier denitrification pathway in MP group at D90. The nitrifier denitrification pathway leads to the transformation of ammonia (NH_3_) into nitrite (NO ^-^) then nitric oxide (NO), nitrous oxide (N_2_O) and finally molecular nitrogen N_2_ (49). Dramatic environmental impacts of NH_3_ are well documented (acidification and eutrophication of ecosystems, biodiversity shifts, ..) and limitation of NH_3_ emission is an important goal to achieve in agricultural industry (50). It has been demonstrated that microaerophilic conditions coupled with low organic matter contents and low pH stop the nitrifier denitrification pathway at the level of N_2_O production, a greenhouse gas with important consequences for environment (49). In our study, bedding samples presented both alkaline pH and high OM values, conditions more prone to promote the nitrifier denitrification pathway, which would thus limit NH_3_ emission and lead to the ultimate transformation into N_2_. Syringate degradation pathways was also observed to be enriched in MP D90. Syringate is a major product in the microbial and chemical degradation of syringyl lignin, with its degradation leading to the production of pyruvate and oxaloacetate (51). Several *Bacillus* strains have been observed among the lignolytic microorganisms identified in plants, soil, wood, or the gut (52), thus the increase of syringate degradation pathway might be linked to the spraying of *Bacillus* through MP application. In addition, two L-arginine degradation pathways producing a biogenic amine (i.e. putrescine) as intermediate metabolite (53,54) were enriched in Control group at D51. Putrescine can be a precursor for other biogenic amines production (spermine, spermidine), and its ingestion has been associated to adverse effects on animal such as decreased DMI and milk production of dairy cows (55), decreased intake, volatile fatty acids production, and total tract DM digestibility in steers (56) or lower growth rate in goat kids (57).

### Use of MP conditioner on RMS-bedding has beneficial impacts on teat skin microbiota

As teats of dairy cows are in direct contact with bedding several hours per day, a focus was made on the impact of MP use on bedding on the bacterial population of teat skin. Teat skin bacterial populations were affected by time, a result previously observed in other studies (58,59). Noteworthy, Hohmann and colleagues observed that an increased frequency of bedding cleaning was associated with lower pathogens load on teat skin (59). Similar to the observations on bedding, a higher bacterial diversity was observed in teat skins of animals from MP group over the trial and it can be hypothesized that the more diverse population of commensal bacteria on teat skin would limit pathogen colonization and proliferation. This observation is strengthened by differential analysis which identified a higher number of several potential mastitis pathogens in Control than in MP group. At the start of the trial, teat skin of Control animals was characterized by presence of LAB (*Lactococcus*, *Latilactobacillus*, *Secundilactobacillus*). Among them, *Lactococcus* genus have been previously identified in bovine mastitis (60). A genus member from Corynebacteriaceae family was observed in higher relative abundance in Control group D01, and among this family, *Corynebacterium* is a recognized genus involved in mastitis cases (61,62). At D90, almost all samples from Control group harbored one taxa from the Staphylococcaeceae family and one assigned to *Truperella*. *Truperella* is opportunistic pathogen causing important diseases in domestic animals, and *T. pyogenes* was isolated in approximately 25 % of the mastitic cows quarters over a 2 years-period (63). Only 2 potential mastitis pathogens were associated to teat skin of MP group : *Acinetobacter* and *Pseudomonas*. Noteworthy the prevalence of *Acinetobacter* was higher in MP but the relative abundance was lower compared to Control group. *Pseudomonas* has been observed in some bovine mastitis (64). However, these potential pathogens have been previously observed on teat skin from healthy cows as well (58,65) and specific qPCR or shotgun sequencing analyses would have been required to detect virulence factors linked to mastitis and thus confirm the lower pathogenic load in one of the experimental group.

### MP bacteria are not disseminated into environment and have no effect on milk yield or composition

Surprisingly, the sprayed bacteria from MP product (*B. velezensis*, *P. acidilactici* and *P. pentosaceus*) were not significantly associated with MP beddings and 2 *Bacillus* ASVs were even associated to Control group. However, due to high phylogenetic proximity among *Bacillus* species, ASVs were assigned only at genus level, leading to an impossible discrimination and tracking of the commercial *Bacillus* strain used in this study. Similarly, *Pediococcus* ASV was identified at genus level and was observed only in MP group D51 at very low relative abundance (0.019 %). As *Pediococcus* was not detected at D01, it seems that the genus was not part of the initial RMS bacterial population. When the application rate of MP decreased (from D31 to D90), *Pediococcus* was not observed anymore in MP group, suggesting a clearing effect and no perennial establishment of this species in the bedding environment and confirming thus the interest of a regular product application.

The absence of *Bacillus* transfer to teat skin and to milk was carefully assessed as this spore former genus is of great concern in dairy industry, being responsible for product defects (34). As previously observed in bedding samples, no *Bacillus* ASV was significantly associated to teat skin samples from MP group, while the relative abundance was higher in Control than MP group (on average 0.29 % and 0.18 % respectively). *Pediococcus* was detected in very low relative abundance only in teat skin samples from MP D51.

No difference in milk yield or milk quality was observed between the 2 groups of animals. Milk total flora and aerobic spore counts were on average 4.25 ± 3.53 log_10_ CFU/ml and 12.94 ± 0.85 spores/ml respectively over the trial. These values are in lines with literature (66–68) and confirm the absence of negative impact of MP utilization on RMS-bedding on milk. The commercial bacteria applied to RMS-bedding were thus neither transferred to teat skin nor to milk over the 3 months trial.

### Use of MP conditioner on RMS-bedding increased cow’s health parameters

Indirect beneficial effects of MP conditioner use on RMS-bedding were observed on several animal health parameters. Soiled udder being one of the main factors triggering mastitis development, a specific attention was given to animal cleanliness. After 51 days of MP utilization on bedding, the animals were characterized by a better hygiene (cleanliness score) and a lower load of bacteria on teat skin, probably linked to the higher DM observed on MP-bedding. This was also reflecting on SCC in milk. In clinical or sub-clinical mastitis, pathogens invade mammary ducts and proliferate. The inflammatory response results notably in white blood cells infiltration to eliminate the pathogenic bacteria (69). These immune cells are identified as SCC in milk and a threshold of 200 000 cells/ml (i.e. 5.3 log_10_ cell/ml) of milk is commonly accepted to characterize an infected cow (70). In our study, from D44 to D76, SCC in milks from Control group were above this threshold and on average 33.67 % higher (5.61 – 5.49 log_10_ SCC/ml) than in MP group (5.13 – 5 log_10_ SCC/ml), indicating that cows from MP group presented lower activation of their immune system reflecting the lower exposition to mastitis pathogens. Several SCC thresholds are applied by different countries to determine the lowest milk quality acceptable, ranging from 400 000 cells/ml in EU to 750 000 cells/ml in the US for bulk tank milk (71–73). High SCC milks have been shown to adversely affect cheese production or pasteurized milk shelf life (74). More specifically in France, a threshold of 250 000 cells/ml has been set above which financial penalties are applied for the farmer (75). In this study, significantly more animals presented milk samples above this limit in Control than in MP group, highlighting the potential economical interest of controlling bedding quality and its further potential benefits on animal health. Mastitis pathogens can be classified into contagious or environmental categories with contagious bacterial species such as *Streptococcus agalactiae* and *Staphylococcus aureus* being spread from an infected cow to a healthy one, usually at milking time (76). Environmental bacteria (*E. coli*, *Klebsiella*, environmental Streptococci, …) come from the cow environment and contrary to contagious mastitis pathogens, it is not possible to totally eliminate them due to their endemic nature. Management practices aiming to improve cleanliness of the animals and their surroundings are the best strategies to control environmental mastitis outbreaks (5,76). Over the trial, the number of effective environmental mastitis pathogens identification was higher in Control group indicating that the use of MP on RMS-bedding led to a better animal hygiene and health and subsequently decrease the potential risk of mastitis onset.

## Conclusion

The use of MP has shown to improve both physico-chemical and bacterial quality of RMS-beddings over a 3-months trial. MP bedding samples were significantly less associated with potential mastitis bacteria. This greater sanitary quality had further impact on animal health as teat skin of animals in MP group was less soiled and presented an improved bacterial composition with less potential mastitis pathogens than in Control animals. Finally, milk quality was not affected by MP application onto RMS-bedding, confirming the absence of bacterial transfer from the commercial product to the milk. Further research is still needed to understand the mechanisms behind the modifications of RMS-bedding bacterial composition observed in this study, and the respective roles of each component of the MP product.

## Material and methods

### Animal Housing

The study was conducted on a commercial farm in Czech Republic. The building was divided in 8 pens with 4 pens (2 on the north and 2 on the south side) dedicated to the trial with an equal repartition between pens submitted to Control or MP treatment. Each pen was composed of 68 cubicles of 2.5 m² and one milking robot (LELY Holding N.V., Maassluis, Netherlands). Bedding in the cubicles was composed of Recycled Manure Solids (RMS) previously prepared from fresh manure from the same farm. Once a week, about 22kg of fresh RMS was added per cubicle and compacted mechanically by tractor passage. Twice a day, the back part of the cubicles was manually scraped to remove organic waste. This experiment was conducted between the 31^st^ of May and the 1^st^ of September 2021. Temperature of the building farm was ranging between 12.7°C ± 2.9 and 20.2°C ± 4.

### Animal selection

A total of 228 Holstein and Simmental crossbreed Holstein cows were enrolled in Control (n=115) and MP (n=113) groups, with parity ranging from 1 to 8 and with DIM ranging from 0 to 853.

### Bedding experimental treatments

In the treated group (MP), the bedding conditioner MANURE PRO (Lallemand SAS, Blagnac, France), composed of live *Bacillus velezensis*, *Pediococcus acidilactici* and *P. pentosaceus* at 5 x 10^9^ CFU/g and cellulolytic enzymes, was sprayed with a backpack sprayer once a week at a concentration of 1g/m²/week for 1 month in the pen, cubicles, exercise area and corridor. During the 2 following months, application rate was decreased to 0.5g/m²/week as the 1-month set-up phase was complete. The application rate from D01 to D30 was voluntary set up at twice the commercial dosage used afterwards in order to prime the effect of the sprayed inoculum on bedding environment. Building surfaces of Control group didn’t receive any treatment.

### Sampling procedures

RMS, bedding and teat skin sampling were done before the first application, at the middle and end of the trial, respectively Day 01, Day 51, and Day 90. The bedding samples were carefully taken with clean gloves at 5cm of depth in 4 locations in the cubicle (2 front, 2 back), avoiding collecting feces. Bedding samples were mixed thoroughly and stored at -80°C until processing. Two replicates were collected per row of cubicles, in 3 rows per pen, for a total of 6 bedding samples per pen per time. Teat skin was sampled using sterile swabs, pre-soaked with AMIES medium (FLMedical, Torreglia, Italy), by gently rubbing 2 opposite teats of the animal. Swabs were kept in buffer according to manufacturer recommendations and stored at -80°C. Teat skin samples were collected on 10 animals per pen (20 per experimental group) balanced in terms of rank of lactation (parity 2 and 3) and with DIM ranging from 70 to 171.

### Sample analysis

#### ○ Bedding

pH values of bedding samples were measured with a pH meter from 10g of fresh bedding diluted into 50ml of CO_2_-free distilled water. DM and OM were analyzed according to the Unified Work Procedures issued by the Central Institute for Supervising and Testing in Agriculture (Czech Republic) based on Commission Regulation (EC) no, 152/2009 (77). Briefly, 50g of fresh bedding sample were dried for 4h at 103°C before weighting for DM calculation. OM was analyzed by ashing 1g of dried samples at 550°C. The measurements were done by the laboratory S.O.S. Skalice n. Svit., s.r.o., Czech Republic.

#### ○ Milk

Milk production was followed every day for each cow individually. Milk conductivity (mS/cm) was measured automatically by milking robots at each milking. Somatic cells count (SCC) was individually followed every 15 days (6 measures over the experimental period) by flow cytometry using Combi Foss (Foss Electric, Hillerød, Denmark).

#### ○ Microbiology

Swabs samples from teats were thoroughly mixed into their respective buffer and an aliquot was used for total flora numeration. Milk total culturable flora was determined by plating an aliquot of raw milk from 20 cows per group. Aerobic spores were enumerated by heating 1ml of milk at 80°C for 10min before enumeration. All microbiology analyzes were done according to ISO 4833-1:2013 (78) at the State Veterinary Institute Jihlava, Czech Republic.

#### ○ Mastitis detection

Based on conductivity measurements provided by Lely robot at each milking, any cow milk above 90 mS/cm was classified under “mastitis suspicion” category while cows below the threshold were classified as “healthy”. Milk samples from “mastitis suspicion” category were sampled and tested through MicroMastTM system (MicroMast, Světlá nad Sázavou, Czech Republic) according to manufacturer recommendations. This on-farm method of mastitis diagnostics allowed the identification of pathogens in milk samples among the main gram negative and positive bacteria responsible for mastitis.

#### ○ Cleanliness score

Cleanliness score was assessed by a trained staff following the grid provided by NZ dairy organization (www.dairynz.co.nz). A score ranging from 0 (clean) to 2 (soiled) was assigned to the back, flank, tail, to the hind leg and to the udder of each animal for a maximum of 6 points being the worst hygienic condition.

### Statistical analysis

Data were analyzed using GraphPad 10.0.0. Microbiological data and SCC were log_10_-transformed before statistical analysis. Evolution of bedding parameters was analyzed through non parametric Mann-Whitney test considering the evolution between D01 and D51 for Period 1, and between D51 and D90 for Period 2. Mixed model with repetition considering Group and Time as fixed factors and the effect of their interaction was applied on animal hygiene parameters, SCC and milk production. Milk production was averaged for each animal per month of treatment application to follow the difference in MP application rate with Month 1 from D01 to D30 at 1g/m²; Month 2 from D31 to D60 and Month 3 from D61 to D90, both at 0.5g/m². Fischer test was applied on suspicious mastitis cases observed in both groups per week and over the trial, as well as on the number of animal presenting SCC in milk above 250 000 cells/ml. Number of suspicious mastitis leading to effective pathogen identification through on-farm plating in each group were compared through non parametric Mann-Whitney test. Mann-Whitney test was also applied to total milk flora and milk aerobic spores data. For all statistical analysis, significant difference was declared at p < 0.05.

### Molecular analysis

#### ○ DNA extraction

DNA extraction was performed from 4g of frozen bedding samples (-80°) or from teat skin swabs with the Quick-DNA™ Fecal/Soil Microbe 96 kit (Zymo Research, Irvine, CA, USA) according to manufacturer’s instructions. Bedding samples and swabs were blended with 20mL of PBS in stomacher bags with 280µM filter, for 1min and 5min respectively, using a stomacher (Smacher, bioMérieux, normal speed). For bedding samples, 3mL of suspensions were first centrifuged at 500rpm for 30s, supernatant was collected and centrifuged at 10000g for 1min. For swab samples, suspensions were centrifuge at 4122g for 30min. Pellets were collected and stored at -20°C until DNA extraction. DNA yield and quality was determined with Nanodrop 1000. DNA extracts were stored at -20 °C until analysis.

#### ○ Amplicon sequencing analysis

Bacterial diversity and taxonomic composition were analysed by 16S rRNA gene amplicon sequencing. The hypervariable V3-V4 regions of the 16S rRNA gene were targeted using the primers set 341F 5’-CCTACGGGAGGCAGCAG-3’ and 806R 5’-GGACTACNVGGGTWTCTAAT-3’. A total of 194 samples were sequenced on a MiSeq high-throughput sequencer (Illumina, San Diego, CA) at the GeT-PlaGe facility (INRAE, Genotoul, Toulouse, France). MiSeq Reagent Kit v3 was used according to the manufacturer’s instruction (Illumina Inc., San Diego, CA) and paired-end 250 bp sequences were generated. Bacterial taxonomic assignment was performed using the DADA2 pipeline 1.22.0 (78) for the pipeline’s steps of filtering, trimming (R1 trimmed at 235bp, R2 trimmed at 220bp), dereplication, sample composition inference and chimera removal. The Silva nr v.138.1 database was used to assign taxonomy from Kingdom to Genus with a minimal boostrap of 30. Diversity analyses were performed with the Phyloseq R package (80). Sequencing run produced a total of 4 462 388 reads used as input for bioinformatic analysis and a total of 3 218 612 reads were kept after trimming, chimeral removal and filtration steps. Rarefaction curves were analyzed to confirm the correct depth sequencing of each sample.

#### ○ Sequencing data analysis

Raw ASV abundances were used for the composition and alpha-diversity analyses. Beta diversity analyses were performed using transformed abundance table with the DESeq2’s variance stabilizing transformation (81). The overall dissimilarity of the microbial community across days and groups was evaluated by principal coordinates analysis (PCoA) based on unweighted Bray-Curtis dissimilarity. The significance of differences between experimental groups was tested by analysis of similarity (PERMANOVA adonis test) and multiple comparisons using Benjamini and Hochberg correction. Indicator species in bedding and on teat skin were identified using the “Indicspecies” package in R (82,83). This method identifies species that are specific to one group with high fidelity (most samples in that group have the species). For this, a genus-level identity table was used as input. Each genus ecological niche preference (Control or MP at each sampling day) was identified using the Pearson’s phi coefficient of association (corrected for unequal sample sizes) using the “Indicspecies” package and 999 permutations. All samples were considered as independent. LEFSe (Linear Discriminant analysis Effet Size) analysis was performed on bedding and teat skin samples using RLE (Relative Log Expression) normalization with a 2.5 LDA score cutoff on identity tables agglomerated down to genus level and a minimum of sample per group set at 5. Functional inference analysis was performed on 16S bedding data following PICRUSt2 pipeline v2.5.0 (84) and HMMER (http://hmmer.org), EPA-NG (85), gappa (86) and SEPP (87) tools for phylogenetic placement of reads, castor tool for hidden state prediction (88) and MinPath tool for pathway inference (89). Statistical analysis was then done using STAMP 2.3.1 software with two groups comparison through two-sided Welch t-test and significance was declared at p < 0.05.

## Declarations

### ● Ethics approval and consent to participate

Not applicable

### ● Consent for publication

Not applicable

### ● Availability of data and material

The sequence datasets generated during the current study are available in the SRA repository, (https://www.ncbi.nlm.nih.gov/sra/) as PRJNA1031159. All the other data are included in this published article and its supplementary information files.

### ● Authors’ contributions

JP and BF designed and supervised the project with contributions from EC. CA performed the DNA extraction. LD performed sequence data analyses. BF and LD performed the statistical analyses. LD wrote the manuscript and prepared the figures. All the authors read and approved the final manuscript.

## ● Acknowledgements

The authors would like to thank the staff from Probionic s.r.o. farm their help in the animal experiment, and Dr Chaucheyras-Durand for her valuable help in reviewing this manuscript.

## Supplementary Figures and Tables

**Supplementary Figure 1:**
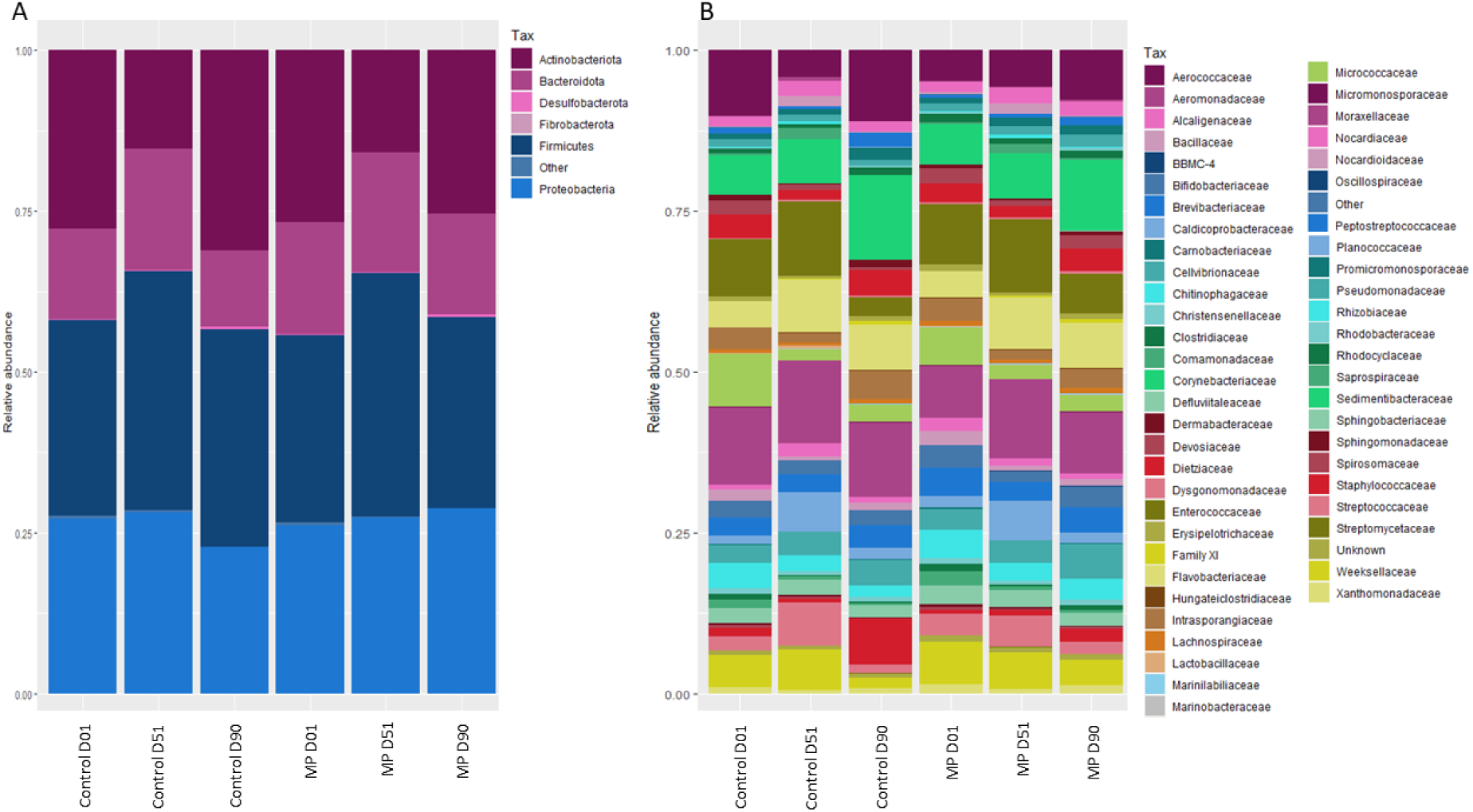
Bacterial composition at phylum level (A) and Family level (B) of Control and MP treated beddings (n=12) collected at D01, D51 and D90.

**Supplementary Figure 2:**
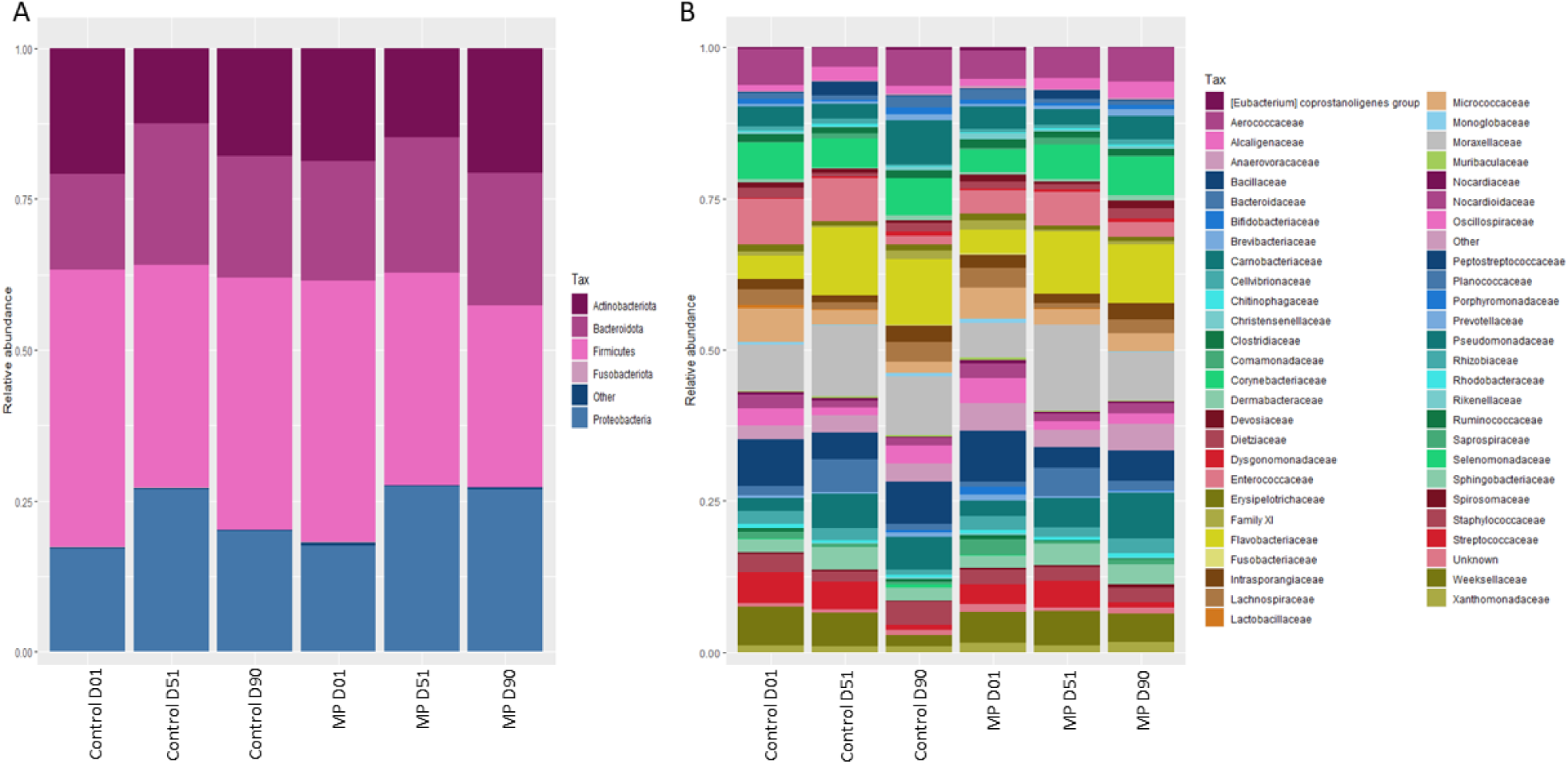
Bacterial composition at A) Phylum and B) Family level of Control and MP treated teat skin samples (n=18) collected at D01, D51 and D90.

**Supplementary Figure 3:**
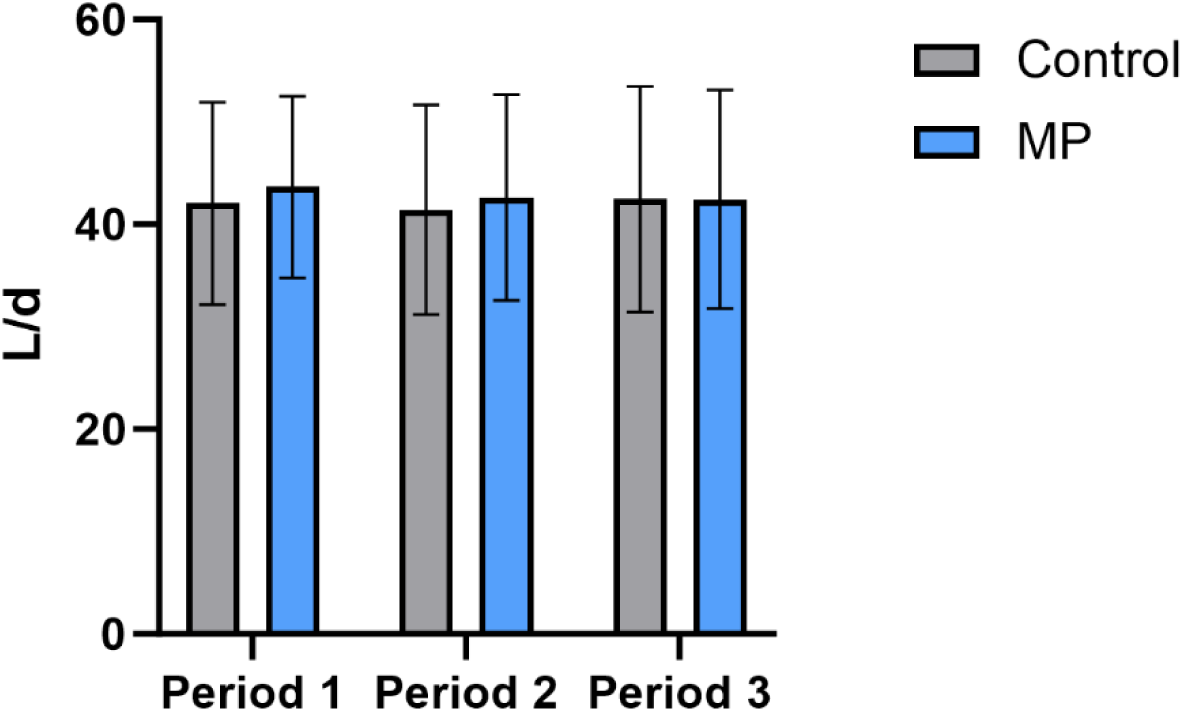
Average milk yield (L/d) for cows in Control (grey, n=113) or MP (blue, n=115) groups in Period 1 (D01-D30), Period 2 (D31-D60) and Period 3 (D61-D90).

**Supplementary Table 1:**
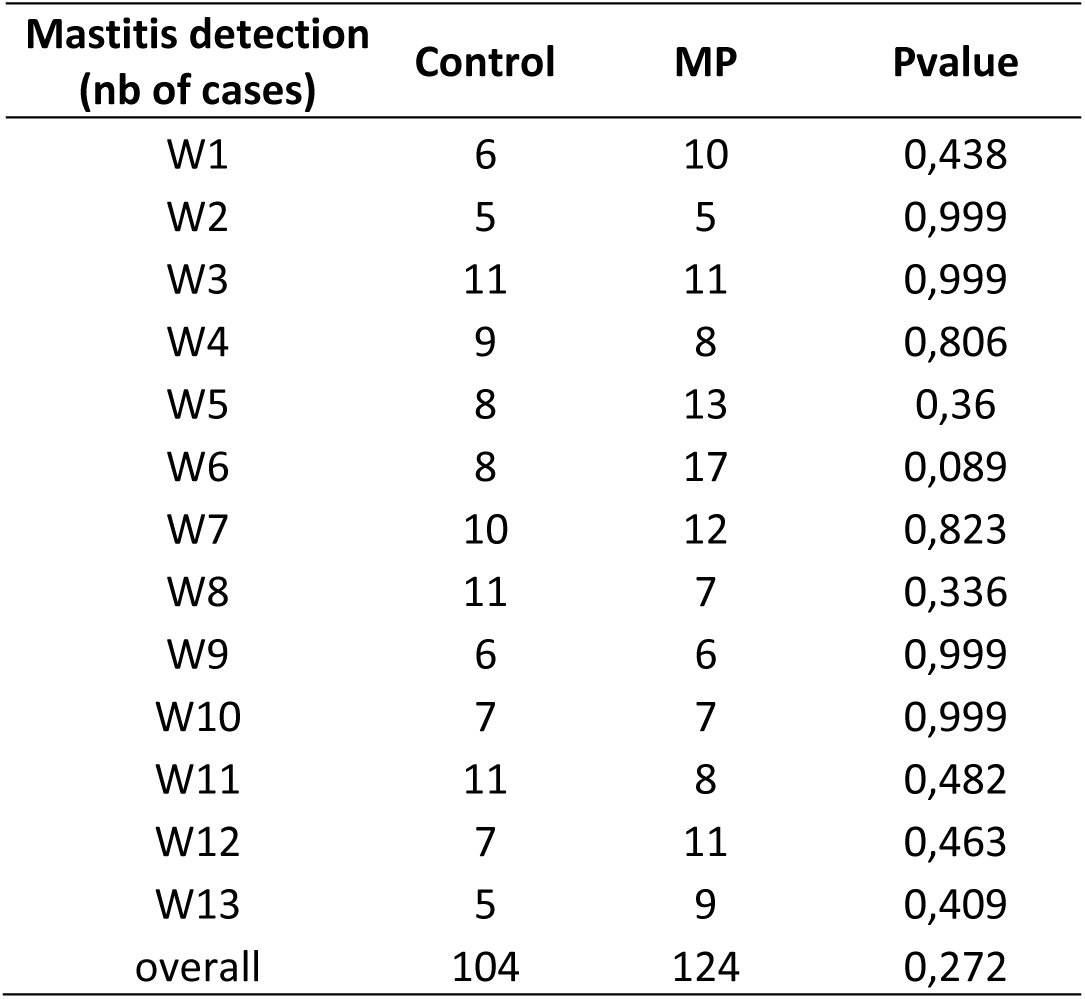
Number of suspicious mastitis cases through milk conductivity in Control (n=113) or MP (n=115) groups by week or over the whole experimental period and Pvalues associated.

| Mastitis detection<br>(nb of cases) | Control | MP | Pvalue |
| --- | --- | --- | --- |
| W1 | 6 | 10 | 0,438 |
| W2 | 5 | 5 | 0,999 |
| W3 | 11 | 11 | 0,999 |
| W4 | 9 | 8 | 0,806 |
| W5 | 8 | 13 | 0,36 |
| W6 | 8 | 17 | 0,089 |
| W7 | 10 | 12 | 0,823 |
| W8 | 11 | 7 | 0,336 |
| W9 | 6 | 6 | 0,999 |
| W10 | 7 | 7 | 0,999 |
| W11 | 11 | 8 | 0,482 |
| W12 | 7 | 11 | 0,463 |
| W13 | 5 | 9 | 0,409 |
| overall | 104 | 124 | 0,272 |

## References

1. Leach KA, Archer SC, Breen JE, Green MJ, Ohnstad IC, Tuer S, et al. Recycling manure as cow bedding: Potential benefits and risks for UK dairy farms. The Veterinary Journal. 1 nov 2015;206(2):123-30.

2. Beauchemin J, Fréchette A, Thériault W, Dufour S, Fravalo P, Thibodeau A. Comparison of microbiota of recycled manure solids and straw bedding used in dairy farms in eastern Canada. Journal of Dairy Science. 1 janv 2022;105(1):389-408.

3. Zdanowicz M, Shelford JA, Tucker CB, Weary DM, von Keyserlingk MAG. Bacterial Populations on Teat Ends of Dairy Cows Housed in Free Stalls and Bedded with Either Sand or Sawdust. Journal of Dairy Science. 1 juin 2004;87(6):1694-701.

4. Vasseur E, Rushen J, Haley DB, de Passillé AM. Sampling cows to assess lying time for on-farm animal welfare assessment. Journal of Dairy Science. 1 sept 2012;95(9):4968-77.

5. Hogan J, Smith KL. Managing Environmental Mastitis. Veterinary Clinics: Food Animal Practice. 1 juill 2012;28(2):217-24.

6. DeVries TJ, Aarnoudse MG, Barkema HW, Leslie KE, von Keyserlingk MAG. Associations of dairy cow behavior, barn hygiene, cow hygiene, and risk of elevated somatic cell count. Journal of Dairy Science. 1 oct 2012;95(10):5730-9.

7. McDougall S, Parker KI, Heuer C, Compton CWR. A review of prevention and control of heifer mastitis via non-antibiotic strategies. Veterinary Microbiology. 16 févr 2009;134(1):177-85.

8. De Vliegher S, Ohnstad I, Piepers S. Management and prevention of mastitis: A multifactorial approach with a focus on milking, bedding and data-management. Journal of Integrative Agriculture. 1 juin 2018;17(6):1214-33.

9. Nonnemann B, Lyhs U, Svennesen L, Kristensen KA, Klaas IC, Pedersen K. Bovine mastitis bacteria resolved by MALDI-TOF mass spectrometry. Journal of Dairy Science. 1 mars 2019;102(3):2515-24.

10. Pang M, Xie X, Bao H, Sun L, He T, Zhao H, et al. Insights Into the Bovine Milk Microbiota in Dairy Farms With Different Incidence Rates of Subclinical Mastitis. Front Microbiol. 16 oct 2018;9:2379.

11. Claxton PD, Ryan D. Bovine mastitis bacteriology. In: Australian Bureau of Animal health. Australia; 1980. p. 8. (Australian Standard Diagnostic Techniques for Animal Diseases; vol. Standing commitee on agriculture and resource management).

12. Francoz D, Wellemans V, Dupré JP, Roy JP, Labelle F, Lacasse P, et al. Invited review: A systematic review and qualitative analysis of treatments other than conventional antimicrobials for clinical mastitis in dairy cows. J Dairy Sci. oct 2017;100(10):7751-70.

13. Seegers H, Fourichon C, Beaudeau F. Production effects related to mastitis and mastitis economics in dairy cattle herds. Veterinary Research. 2003;34(5):475-91.

14. Schepers JA, Dijkhuizen AA. The economics of mastitis and mastitis control in dairy cattle: a critical analysis of estimates published since 1970. Preventive Veterinary Medicine. 1 mars 1991;10(3):213-24.

15. Halasa T, Huijps K, Østerås O, Hogeveen H. Economic effects of bovine mastitis and mastitis management: A review. Veterinary Quarterly. 1 janv 2007;29(1):18-31.

16. Ruegg PL. A 100-Year Review: Mastitis detection, management, and prevention. Journal of Dairy Science. 1 déc 2017;100(12):10381-97.

17. Gonçalves JL, de Campos JL, Steinberger AJ, Safdar N, Kates A, Sethi A, et al. Incidence and Treatments of Bovine Mastitis and Other Diseases on 37 Dairy Farms in Wisconsin. Pathogens. nov 2022;11(11):1282.

18. Krishnamoorthy P, Goudar AL, Suresh KP, Roy P. Global and countrywide prevalence of subclinical and clinical mastitis in dairy cattle and buffaloes by systematic review and meta-analysis. Research in Veterinary Science. 1 mai 2021;136: 561-86.

19. Patel K, Godden SM, Royster E, Crooker BA, Timmerman J, Fox L. Relationships among bedding materials, bedding bacteria counts, udder hygiene, milk quality, and udder health in US dairy herds. Journal of Dairy Science. 1 nov 2019;102(11):10213-34.

20. Rowbotham RF, Ruegg PL. Associations of selected bedding types with incidence rates of subclinical and clinical mastitis in primiparous Holstein dairy cows. Journal of Dairy Science. 1 juin 2016;99(6):4707-17.

21. Fréchette A, Fecteau G, Côté C, Dufour S. Association Between Recycled Manure Solids Bedding and Subclinical Mastitis Incidence: A Canadian Cohort Study. Frontiers in Veterinary Science [Internet]. 2022 [cité 9 août 2023];9. Disponible sur: https://www.frontiersin.org/articles/10.3389/fvets.2022.859858

22. Robles I, Kelton DF, Barkema HW, Keefe GP, Roy JP, von Keyserlingk M a. G, et al. Bacterial concentrations in bedding and their association with dairy cow hygiene and milk quality. Animal. mai 2020;14(5):1052-66.

23. Alanis VM, Zurakowski M, Pawloski D, Tomazi T, Nydam DV, Ospina PA. Description of the Characteristics of Five Bedding Materials and Association With Bulk Tank Milk Quality on Five New York Dairy Herds. Frontiers in Veterinary Science [Internet]. 2021 [cité 14 févr 2023];8. Disponible sur: https://www.frontiersin.org/articles/10.3389/fvets.2021.636833

24. Paduch JH, Mohr E, Krömker V. The association between bedding material and the bacterial counts of Staphylococcus aureus, Streptococcus uberis and coliform bacteria on teat skin and in teat canals in lactating dairy cattle. Journal of Dairy Research. mai 2013;80(2):159-64.

25. Proietto RL, Hinckley LS, Fox LK, Andrew SM. Evaluation of a clay-based acidic bedding conditioner for dairy cattle bedding. Journal of Dairy Science. 1 févr 2013;96(2):1044-53.

26. Hogan JS, Wolf SL, Petersson-Wolfe CS. Bacterial Counts in Organic Materials Used as Free-Stall Bedding Following Treatment with a Commercial Conditioner. Journal of Dairy Science. 1 févr 2007;90(2):1058-62.

27. Hogan JS, Bogacz VL, Thompson LM, Romig S, Schoenberger PS, Weiss WP, et al. Bacterial counts associated with sawdust and recycled manure bedding treated with commercial conditioners. J Dairy Sci. août 1999;82(8):1690-5.

28. Muck RE, Nadeau EMG, McAllister TA, Contreras-Govea FE, Santos MC, Kung L. Silage review: Recent advances and future uses of silage additives. Journal of Dairy Science. 1 mai 2018;101(5):3980-4000.

29. Greff B, Szigeti J, Nagy Á, Lakatos E, Varga L. Influence of microbial inoculants on co-composting of lignocellulosic crop residues with farm animal manure: A review. Journal of Environmental Management. 15 janv 2022;302:114088.

30. Ayilara MS, Babalola OO. Bioremediation of environmental wastes: the role of microorganisms. Frontiers in Agronomy [Internet]. 2023 [cité 4 sept 2023];5. Disponible sur: https://www.frontiersin.org/articles/10.3389/fagro.2023.1183691

31. Pang X, Song X, Chen M, Tian S, Lu Z, Sun J, et al. Combating biofilms of foodborne pathogens with bacteriocins by lactic acid bacteria in the food industry. Comprehensive Reviews in Food Science and Food Safety. 2022;21(2):1657-76.

32. Guéneau V, Rodiles A, Frayssinet B, Piard JC, Castex M, Plateau-Gonthier J, et al. Positive biofilms to control surface-associated microbial communities in a broiler chicken production system -a field study. Frontiers in Microbiology [Internet]. 2022 [cité 4 sept 2023];13. Disponible sur: https://www.frontiersin.org/articles/10.3389/fmicb.2022.981747

33. Adeniji AA, Loots DT, Babalola OO. Bacillus velezensis: phylogeny, useful applications, and avenues for exploitation. Appl Microbiol Biotechnol. 1 mai 2019;103(9):3669-82.

34. Heyndrickx M, Scheldeman P. Bacilli associated with spoilage in dairy products and other food. In: Applications and systematics of Bacillus and relatives [Internet]. John Wiley & Sons, Ltd; 2002 [cité 3 janv 2023]. p. 64-82. Disponible sur: https://onlinelibrary.wiley.com/doi/abs/10.1002/9780470696743.ch6

35. Ray T, Gaire TN, Dean CJ, Rowe S, Godden SM, Noyes NR. The microbiome of common bedding materials before and after use on commercial dairy farms. Animal Microbiome. 7 mars 2022;4(1):18.

36. European Union. REGULATION (EC) No 1069/2009 OF THE EUROPEAN PARLIAMENT AND OF THE COUNCIL of 21 October 2009 laying down health rules as regards animal by-products and derived products not intended for human consumption and repealing Regulation (EC) No 1774/2002 (Animal by-products Regulation) [Internet]. 2009 p. 1-33. Report No.: 2009/1069/EC. Disponible sur: chrome-extension://efaidnbmnnnibpcajpcglclefindmkaj/https://eur-lex.europa.eu/LexUriServ/LexUriServ.do?uri=OJ:L:2009:300:0001:0033:en:PDF

37. Bradley AJ, Leach KA, Green MJ, Gibbons J, Ohnstad IC, Black DH, et al. The impact of dairy cows’ bedding material and its microbial content on the quality and safety of milk – A cross sectional study of UK farms. International Journal of Food Microbiology. 23 mars 2018;269:36-45.

38. Costa-Roura S, Villalba D, Balcells J, De la Fuente G. First Steps into Ruminal Microbiota Robustness. Animals. janv 2022;12(18):2366.

39. Weimer PJ. Redundancy, resilience, and host specificity of the ruminal microbiota: implications for engineering improved ruminal fermentations. Front Microbiol. 10 avr 2015;6:296.

40. Gagliardi A, Totino V, Cacciotti F, Iebba V, Neroni B, Bonfiglio G, et al. Rebuilding the Gut Microbiota Ecosystem. Int J Environ Res Public Health. août 2018;15(8):1679.

41. Song X, Huang X, Xu H, Zhang C, Chen S, Liu F, et al. The prevalence of pathogens causing bovine mastitis and their associated risk factors in 15 large dairy farms in China: An observational study. Veterinary Microbiology. 1 août 2020;247:108757.

42. Różańska H, Lewtak-Piłat A, Kubajka M, Weiner M. Occurrence of Enterococci in Mastitic Cow’s Milk and their Antimicrobial Resistance. J Vet Res. 22 mars 2019;63(1):93-7.

43. Kim HJ, Youn HY, Kang HJ, Moon JS, Jang YS, Song KY, et al. Prevalence and Virulence Characteristics of Enterococcus faecalis and Enterococcus faecium in Bovine Mastitis Milk Compared to Bovine Normal Raw Milk in South Korea. Animals (Basel). 30 mai 2022;12(11):1407.

44. Nagaraja TG, Narayanan SK, Stewart GC, Chengappa MM. Fusobacterium necrophorum infections in animals: Pathogenesis and pathogenic mechanisms. Anaerobe. 1 août 2005;11(4):239-46.

45. Jousimies-Somer H, Pyörälä S, Kanervo A. Susceptibilities of bovine summer mastitis bacteria to antimicrobial agents. Antimicrobial Agents and Chemotherapy. janv 1996;40(1):157-60.

46. Madsen M, Høi Sørensen G, Aalbaek B. Summer mastitis in heifers: a bacteriological examination of secretions from clinical cases of summer mastitis in Denmark. Veterinary Microbiology. 1 mai 1990;22(4):319-28.

47. Fabbi M, Pastoris MC, Scanziani E, Magnino S, Di Matteo L. Epidemiological and Environmental Investigations of Legionella pneumophila Infection in Cattle and Case Report of Fatal Pneumonia in a Calf. J Clin Microbiol. juill 1998;36(7):1942-7.

48. Lye D, Fout GS, Crout SR, Danielson R, Thio CL, Paszko-Kolva CM. Survey of ground, surface, and potable waters for the presence of Legionella species by EnviroAmpR PCR Legionella Kit, culture, and immunofluorescent staining. Water Research. 1 févr 1997;31(2):287-93.

49. Wrage N, Velthof GL, van Beusichem ML, Oenema O. Role of nitrifier denitrification in the production of nitrous oxide. Soil Biology and Biochemistry. 1 oct 2001;33(12):1723-32.

50. Wyer KE, Kelleghan DB, Blanes-Vidal V, Schauberger G, Curran TP. Ammonia emissions from agriculture and their contribution to fine particulate matter: A review of implications for human health. Journal of Environmental Management. 1 déc 2022;323:116285.

51. Gao J, Ellis LBM, Wackett LP. The University of Minnesota Biocatalysis/Biodegradation Database: improving public access. Nucleic Acids Research. 1 janv 2010;38(suppl_1):D488-91.

52. Grgas D, Rukavina M, Bešlo D, Štefanac T, Crnek V, Šikić T, et al. The Bacterial Degradation of Lignin—A Review. Water. janv 2023;15(7):1272.

53. MetaCyc EC 4.1.1.17 [Internet]. [cité 27 oct 2023]. Disponible sur: https://biocyc.org/META/NEW-IMAGE?type=REACTION&object=ORNDECARBOX-RXN

54. MetaCyc superpathway of L-arginine, putrescine, and 4-aminobutanoate degradation [Internet]. [cité 27 oct 2023]. Disponible sur: https://biocyc.org/META/NEW-IMAGE?type=PATHWAY&object=ARGDEG-PWY

55. Lingaas F, Tveit B. Etiology of acetonemia in Norwegian cattle. 2. Effect of butyric acid, valeric acid, and putrescine. J Dairy Sci. sept 1992;75(9):2433-9.

56. Phuntsok T, Froetschel MA, Amos HE, Zheng M, Huang YW. Biogenic Amines in Silage, Apparent Postruminal Passage, and the Relationship Between Biogenic Amines and Digestive Function and Intake by Steers. Journal of Dairy Science. 1 août 1998;81(8):2193-203.

57. Fusi E, Rossi L, Rebucci R, Cheli F, Di Giancamillo A, Domeneghini C, et al. Administration of biogenic amines to Saanen kids: effects on growth performance, meat quality and gut histology. Small Ruminant Research. 1 juin 2004;53(1):1-7.

58. Verdier-Metz I, Delbès C, Bouchon M, Pradel P, Theil S, Rifa E, et al. Influence of Post-Milking Treatment on Microbial Diversity on the Cow Teat Skin and in Milk. Dairy. juin 2022;3(2):262-76.

59. Hohmann MF, Wente N, Zhang Y, Krömker V. Bacterial Load of the Teat Apex Skin and Associated Factors at Herd Level. Animals (Basel). 14 sept 2020;10(9):1647.

60. Rodrigues MX, Lima SF, Higgins CH, Canniatti-Brazaca SG, Bicalho RC. The Lactococcus genus as a potential emerging mastitis pathogen group: A report on an outbreak investigation. Journal of Dairy Science. 1 déc 2016;99(12):9864-74.

61. Kittl S, Studer E, Brodard I, Thomann A, Jores J. Corynebacterium uberis sp. nov. frequently isolated from bovine mastitis. Syst Appl Microbiol. juill 2022;45(4):126325.

62. Gonçalves JL, Tomazi T, Barreiro JR, Beuron DC, Arcari MA, Lee SHI, et al. Effects of bovine subclinical mastitis caused by Corynebacterium spp. on somatic cell count, milk yield and composition by comparing contralateral quarters. The Veterinary Journal. 1 mars 2016;209:87-92.

63. Ishiyama D, Mizomoto T, Ueda C, Takagi N, Shimizu N, Matsuura Y, et al. Factors affecting the incidence and outcome of Trueperella pyogenes mastitis in cows. J Vet Med Sci. mars 2017;79(3):626-31.

64. Schauer B, Wald R, Urbantke V, Loncaric I, Baumgartner M. Tracing Mastitis Pathogens—Epidemiological Investigations of a Pseudomonas aeruginosa Mastitis Outbreak in an Austrian Dairy Herd. Animals (Basel). 22 janv 2021;11(2):279.

65. Frétin M, Martin B, Rifa E, Isabelle VM, Pomiès D, Ferlay A, et al. Bacterial community assembly from cow teat skin to ripened cheeses is influenced by grazing systems. Scientific Reports. 9 janv 2018;8(1):200.

66. Verdier-Metz I, Gagne G, Bornes S, Monsallier F, Veisseire P, Delbès-Paus C, et al. Cow Teat Skin, a Potential Source of Diverse Microbial Populations for Cheese Production. Appl Environ Microbiol. 15 janv 2012;78(2):326-33.

67. Cebeci T. A Survey of Raw Milk For Microbiological Quality and Typing of Foodborne Pathogens by MALDI-TOF MS. ADÜ ZİRAAT DERG. 31 déc 2019;16(2):185-91.

68. Ortuzar J, Martinez B, Bianchini A, Stratton J, Rupnow J, Wang B. Quantifying changes in spore-forming bacteria contamination along the milk production chain from farm to packaged pasteurized milk using systematic review and meta-analysis. Food Control. 1 avr 2018;86:319-31.

69. Goulart DB, Mellata M. Escherichia coli Mastitis in Dairy Cattle: Etiology, Diagnosis, and Treatment Challenges. Frontiers in Microbiology [Internet]. 2022 [cité 12 sept 2023];13. Disponible sur: https://www.frontiersin.org/articles/10.3389/fmicb.2022.928346

70. Moroni P, Nydam DV, Ospina PA, Scillieri-Smith JC, Virkler PD, Watters RD, et al. 8 - Diseases of the Teats and Udder. In: Peek SF, Divers TJ, éditeurs. Rebhun’s Diseases of Dairy Cattle (Third Edition) [Internet]. Elsevier; 2018 [cité 21 sept 2023]. p. 389-465. Disponible sur: https://www.sciencedirect.com/science/article/pii/B9780323390552000085

71. U.S. Department of Health and Human Services. Grade « A » pasteurized milk ordinance. (Includes provisions from Condensed and Dry Whe the Grade “A” Condensed and Dry Milk Products and y--Supplement I to the Grade “A” PMO) [Internet]. Food and Drug Administration; 2019 [cité 16 oct 2023] p. 447. (Public health Service). Disponible sur: chrome-extension://efaidnbmnnnibpcajpcglclefindmkaj/https://www.fda.gov/media/140394/download

72. Regulation (EC) No 853/2004 of the European Parliament and of the Council of 29 April 2004 laying down specific hygiene rules for food of animal origin [Internet]. OJ L avr 29, 2004. Disponible sur: http://data.europa.eu/eli/reg/2004/853/oj/eng

73. Règlement (CE) n° 854/2004 du Parlement européen et du Conseil du 29 avril 2004 fixant les règles spécifiques d’organisation des contrôles officiels concernant les produits d’origine animale destinés à la consommation humaine [Internet]. OJ L avr 29, 2004. Disponible sur: http://data.europa.eu/eli/reg/2004/854/oj/fra

74. Moradi M, Omer AK, Razavi R, Valipour S, Guimarães JT. The relationship between milk somatic cell count and cheese production, quality and safety: A review. International Dairy Journal. 1 févr 2021;113:104884.

75. Freiss julien. Evolution de la qualité du lait lors de l’installation d’un robot de traite: description et facteurs de variation. [Internet] [Thèse Vétérinaire]. [Nantes]: Ecole Nationale Vétérinaire de Nantes; 2009. Disponible sur: https://theses.hal.science/tel-00600088

76. Garcia A. Contagious vs. Environmental Mastitis. SDSU Extension Extra Archives [Internet]. 1 janv 2004; Disponible sur: https://openprairie.sdstate.edu/extension_extra/126

77. European Union PO of the E. CELEX1, COMMISSION REGULATION (EC) No 152/2009 of 27 January 2009 laying down the methods of sampling and analysis for the official control of feed (Text with EEA relevance) [Internet]. Bruxelles: Commission européenne; 2012 avr [cité 7 sept 2023] p. 157. Report No.: 02009R0152-20120418. Disponible sur: https://op.europa.eu/fr/publication-detail/-/publication/a72a297b-a867-4424-9f5f-a3c687ba2960/language-fr

78. ISO/TC 34/SC 9 Microbiology. ISO 4833-1:2013 Microbiology of the food chain — Horizontal method for the enumeration of microorganisms — Part 1: Colony count at 30 °C by the pour plate technique [Internet]. 2013 sept [cité 7 sept 2023] p. 9. (07.100.30 Food microbiology). Disponible sur: https://www.iso.org/standard/53728.html

79. Callahan BJ, McMurdie PJ, Rosen MJ, Han AW, Johnson AJA, Holmes SP. DADA2: High-resolution sample inference from Illumina amplicon data. Nature Methods. juill 2016;13(7):581-3.

80. McMurdie PJ, Holmes S. phyloseq: An R Package for Reproducible Interactive Analysis and Graphics of Microbiome Census Data. PLOS ONE. 22 avr 2013;8(4):e61217.

81. Love MI, Huber W, Anders S. Moderated estimation of fold change and dispersion for RNA-seq data with DESeq2. Genome Biology. 5 déc 2014;15(12):550.

82. De Cáceres M, Legendre P. Associations between species and groups of sites: indices and statistical inference. Ecology. déc 2009;90(12):3566-74.

83. Legendre P, De Cáceres M. Beta diversity as the variance of community data: dissimilarity coefficients and partitioning. Ecology Letters. 2013;16(8):951-63.

84. Douglas GM, Maffei VJ, Zaneveld JR, Yurgel SN, Brown JR, Taylor CM, et al. PICRUSt2 for prediction of metagenome functions. Nat Biotechnol. juin 2020;38(6):685-8.

85. Barbera P, Kozlov AM, Czech L, Morel B, Darriba D, Flouri T, et al. EPA-ng: Massively Parallel Evolutionary Placement of Genetic Sequences. Systematic Biology. 1 mars 2019;68(2):365-9.

86. Czech L, Barbera P, Stamatakis A. Genesis and Gappa: processing, analyzing and visualizing phylogenetic (placement) data. Bioinformatics. 15 mai 2020;36(10):3263-5.

87. Mirarab S, Nguyen N, Warnow T. SEPP: SATé-Enabled Phylogenetic Placement. In: Biocomputing 2012 [Internet]. WORLD SCIENTIFIC; 2011 [cité 16 févr 2023]. p. 247-58. Disponible sur: https://www.worldscientific.com/doi/abs/10.1142/9789814366496_0024

88. Louca S, Doebeli M. Efficient comparative phylogenetics on large trees. Bioinformatics. 15 mars 2018;34(6):1053-5.

89. Ye Y, Doak TG. A Parsimony Approach to Biological Pathway Reconstruction/Inference for Genomes and Metagenomes. PLOS Computational Biology. 14 août 2009;5(8):e1000465.

